# A bifunctional RNase H protein supports DNA replication and transcription in nucleus-forming *Chimalliviridae* phages

**DOI:** 10.64898/2026.10.01.755736

**Authors:** Phoolwanti Rani, Niklas Klusch, Shannon M. Barret, Taylor Forman, Makaela Levine, Margot Riggi, Caroline Rees, Rui Liu, Lizbeth Y. Hoffman, Meline Norquist, Dwaipayan Basu, Koe Inlow, Emily G. Armbruster, Chase J. Morgan, Joe Pogliano, Kevin D. Corbett, Elizabeth Villa

## Abstract

Jumbo bacteriophages of the Chimalliviridae family replicate their genomes within a proteinaceous phage nucleus that spatially segregates viral DNA replication and transcription from translation. While the phage nucleus protects viral DNA from host defenses, it also imposes unique constraints on genome replication and gene expression. Here we identify the phage-encoded protein Rhc (RNase H-clamp) as a central mediator of genome transactions within the phage nucleus. Using the *Escherichia coli* infecting bacteriophage Goslar, we show that Rhc functions as an essential component of a non-canonical three-subunit DNA polymerase complex composed of DNAPN, DNAPC, and Rhc. Rhc physically interacts with the DNAPC subunit, and both the catalytic activity of Rhc’s N-terminal RNase H domain and its C-terminal sliding clamp-derived DNA binding domain are required for genome replication, nuclear maturation, and productive infection. Further, Goslar Rhc also supports transcription within the phage nucleus through a function that is separable from its DNA replication function. Rhc depletion results in reduced transcription of a subset of late genes, in a pattern that suggests a role for Rhc in resolving R-loops. Imaging and cryo-electron tomography reveal that loss of Rhc arrests phage development at a nascent nuclear stage, uncoupling nuclear assembly from maturation. Together, these findings identify Rhc as a multifunctional nuclear factor that mediates both DNA replication and transcription in nucleus-forming jumbo phages, revealing how viruses repurpose a conserved enzymatic activity to coordinate gene expression within a compartmentalized replication environment.

## INTRODUCTION

Large-genome bacteriophages (phages) of the *Chimalliviridae* family (here termed chimalliviruses) possess a distinctive life cycle, reorganizing infected bacterial cells by enclosing the phage’s DNA genome within a proteinaceous ‘phage nucleus’ that spatially separates genome replication and transcription from protein translation [1–3]. The phage nucleus and an earlier membrane-bound organelle called the early phage infection (EPI) vesicle serve to protect phage DNA from host-encoded defenses like restriction endonucleases and CRISPR nucleases [4–8]. While the phage nucleus and EPI vesicle provide broad protection against host defenses, they introduce challenges that the phage must overcome to replicate successfully. Because the EPI vesicle and phage nucleus spatially segregate genomic DNA from the host-cell cytoplasm, mRNAs produced in these compartments must be translocated to the cytoplasm for translation by host ribosomes [9,10]. Similarly, in late infections a number of phage- and host-encoded proteins must be specifically imported into the phage nucleus [11,12]. Finally, replicated genomic DNA must be translocated across the phage nuclear shell for packaging into nascent virions. To overcome these challenges, chimalliviruses possess dedicated machinery for nucleic acid and protein transport across the nuclear shell.

Comprehensive bioinformatics analyses have shown that chimalliviruses encode a distinctive, highly diverged repertoire of proteins predicted to play roles in DNA replication, DNA repair, and mRNA transcription [13]. This repertoire includes two multi-subunit RNA polymerases (RNAPs): The “virion” RNAP (vRNAP) is packaged in the phage capsid and injected into a new host along with genomic DNA into the EPI vesicle, and this enzyme is responsible for early-gene transcription. After the genomic DNA has been transferred into the phage nucleus, the “non-virion” RNAP (nvRNAP) is produced and imported into the phage nucleus, and this enzyme is responsible for transcription of late genes [7,14]. Chimalliviruses also encode two proteins with homology to DNA polymerases (DNAPs): one predicted to possess the N-terminal exonuclease domain (here termed DNAP^N^), and the second predicted to possess the C-terminal polymerase domain (here termed DNAP^C^) [13]. It is not known whether these proteins function together or separately. Chimalliviruses also encode a diverged DnaB-like replicative helicase, but notably lack identifiable sliding clamp proteins, which promote processivity of the associated DNA polymerase, or primases, which synthesize RNA primers.

A particularly enigmatic component of chimalliviruses’ DNA replication/repair/transcription toolkit is a conserved, multi-domain RNase H protein (termed Rhc hereafter for RNase H-clamp) [12,15]. The Rhc N-terminal domain resembles RNase HI enzymes that degrade RNA:DNA hybrids, whereas its C-terminal domain lacks obvious sequence homology to characterized proteins. In PhiKZ and PhiPA3, antisense oligonucleotide-mediated knockdown of Rhc strongly reduces DNA replication [15]. Intriguingly, a nuclease-dead Rhc mutant fully rescued DNA replication and phage titer in PhiKZ, suggesting that this protein may serve a structural rather than a catalytic role in DNA replication [15]. A concurrent study finds that PhiKZ Rhc associates transiently with the DNAP^N^-DNAP^C^ complex and contributes to DNA polymerase activity *in vitro* by binding replicated DNA emerging from the enzyme’s active site (Basu *et al*., *bioRxiv*). Still unknown is how significantly Rhc contributes to DNA replication in other chimalliviruses, whether its RNase H catalytic activity contributes to DNA replication, and whether Rhc performs additional nuclear functions independent of DNA replication.

Here, we characterize Rhc’s roles in the *E. coli*-infecting chimallivirus Goslar. We find that in Goslar, late-gene transcription is independent of DNA replication, and that Rhc is separately required for both DNA replication and late-gene transcription. Rhc’s uncharacterized C-terminal domain is structurally related to DNA sliding clamp subunits, and directly associates with the phage-encoded DNA polymerase to boost its activity. In contrast to prior findings in PhiKZ, we find that Goslar Rhc is required for transcription of late genes within the phage nucleus, independent of its association with the DNA polymerase, but dependent on nucleic acid binding by its C-terminal clamp domain. Overall, these data define the structure and functions of a distinctive multidomain RNase H-like protein in chimalliviruses, and reveal how this protein contributes to both DNA replication and transcription during infections.

## RESULTS

### Two domains in Rhc contribute to DNA replication and phage developmental cycle

The *E. coli*-infecting chimallivirus Goslar encodes two proteins annotated as RNase H proteins: gp177 is related to T4 RNase H, a flap endonuclease-related protein, and gp190 (here termed Rhc) is related to RNase HI protein family. We found that gp177 was not required for phage replication (Figure S1) and focused on Rhc. AlphaFold structure predictions of Rhc reveal an N-terminal RNase HI-like domain containing a conserved catalytic site (Figures 1A,B, S2), and a C-terminal domain distantly related to DNA polymerase sliding-clamp proteins including eukaryotic PCNA (proliferating cell nuclear antigen) and the 9-1-1 complex (RAD9, RAD1, HUS1) (Figure 1B, Table S1). We therefore refer to the N-terminal region of Rhc as the RNase H domain, and the C-terminal domain as the clamp domain (Figure 1A).

**Figure 1.**
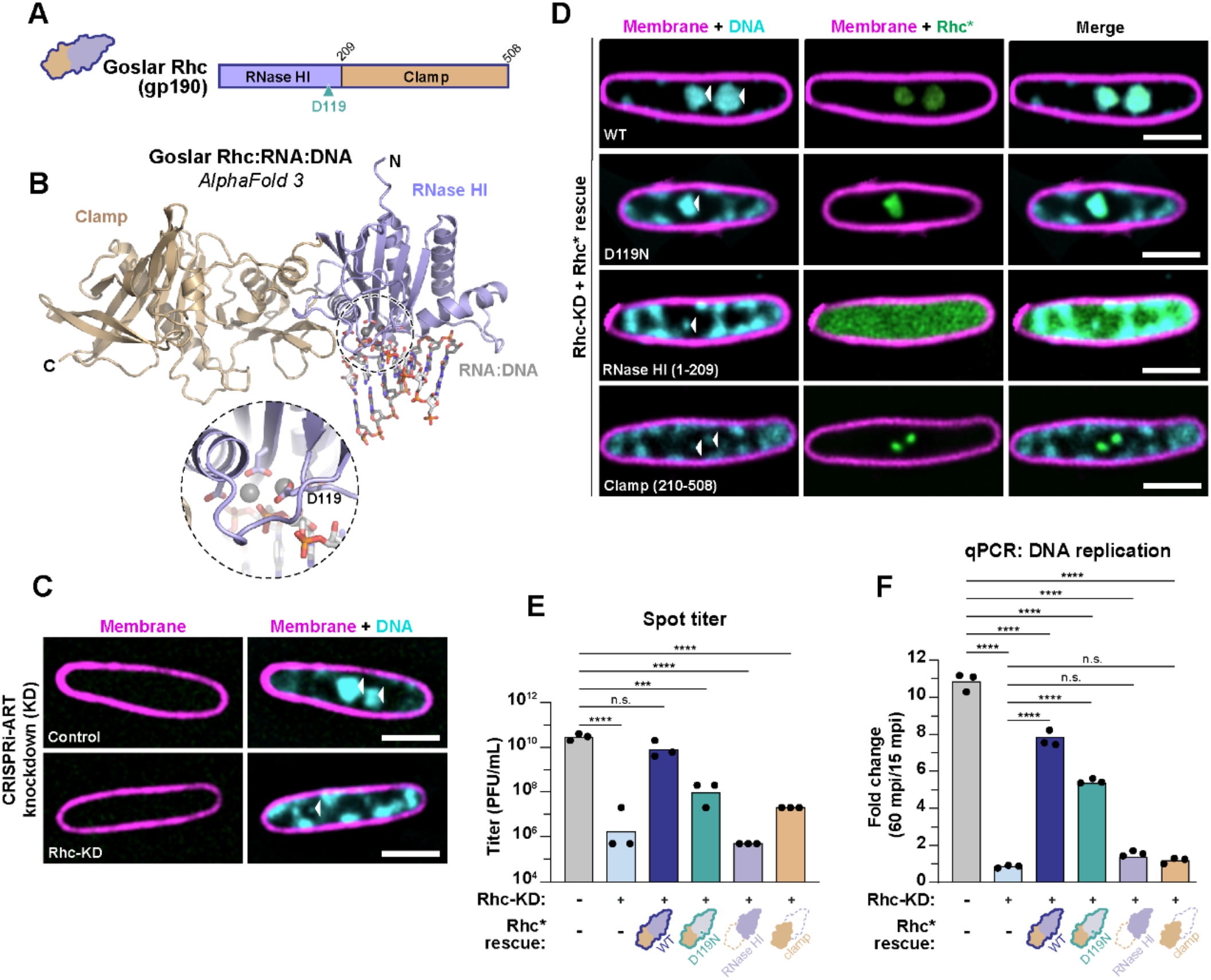
The RNase HI and clamp domains are required for phage nucleus maturation, genome replication, and productive infection. **(A)** Domain organization of Goslar Rhc (gp190) with RNase HI-like and sliding clamp-like domains labeled, and the putative catalytic aspartate residue in each RNase HI-like domain indicated. **(B)** AlphaFold3-predicted structure of the Rhc-RNA:DNA complex. The RNase HI domain is shown in purple, the clamp domain in tan, and the RNA:DNA hybrid as white/gray sticks. *Inset:* Enlarged view of the predicted Rhc catalytic site, highlighting the putative catalytic residue D119. Two predicted Mg2+ ions are shown as gray spheres. See **Figure S2** for details. **(C)** Representative fluorescence microscopy images of Goslar-infected *E. coli* cells with CRISPRi-ART knockdowns. NT: non-targeting control; Rhc-KD: Goslar gp190 knockdown (see **Figure S3** for experimental design) Outer membranes (FM4-64) are shown in magenta and DNA (DAPI) in cyan. Arrowheads indicate phage nuclear DNA. **(D)** Complementation of CRISPRi-ART Rhc knockdown with knockdown-resistant Rhc (Rhc*) and Variants (D119N, RNase HI, Clamp). Outer membranes (FM4-64) are shown in magenta, DNA (DAPI) in cyan, and Rhc in green. WT: wild-type. Arrowheads indicate phage nuclear DNA. **(E)** Spot-titer analysis of infectious phage production under the indicated knockdown and complementation conditions. Titers are reported as log_10_ PFU/mL. Individual biological replicates are shown as black dots, and bars indicate the mean. Statistical significance was determined using the indicated comparisons. ns, not significant; ***P < 0.001; ****P < 0.0001. **(F)** qPCR analysis of phage DNA replication, expressed as the fold change in phage DNA abundance at 60 minutes post-infection relative to 15 minutes post-infection. Bars represent the mean ± s.e.m.; black dots indicate three biological replicates. Statistical significance was determined by one-way ANOVA with Šídák’s multiple-comparisons test. ns, not significant; ****P < 0.0001. Scale bars: (C,D) 2μm.

To determine the functions of Rhc during Goslar infection, we first blocked its synthesis using CRISPRi-ART [16] (Figure S3A) and examined infection progression by fluorescence microscopy. Rhc knockdown resulted in a failure to develop a large DNA-containing phage nucleus (Figure 1C). Phage nuclear growth was rescued by expression of a CRISPRi-ART-resistant complementation vector expressing recoded Rhc (Rhc*) fused to GFP (Figure 1D), demonstrating that the nuclear-growth defect results specifically from loss of Rhc, rather than off-target effects of the knockdown. To test the roles of Rhc’s two domains, we expressed several CRISPRi-ART resistant Rhc* GFP-labelled constructs: a D119N RNase H catalytic-dead full-length mutant (D119N), the isolated N-terminal RNase H domain (residues 1-209, RNase HI), and the isolated C-terminal clamp domain (residues 210-508, clamp). Expression of the Rhc* D119N mutant largely rescued phage nuclear growth, while the isolated RNase HI and clamp domains did not (Figure 1D). The isolated N-terminal RNase H domain failed to localize to the nascent phage nucleus (Figures 1D, S4), while the clamp domain localized robustly to nascent phage nuclei (Figure 1D), consistent with earlier observations of PhiKZ Rhc localization [12].

Next, we tested Goslar phage titer and directly measured DNA replication using qPCR for the Rhc knockdown and all Rhc* constructs. Rhc knockdown caused a ~4-log reduction in Goslar phage titer (Figure 1E), and an almost complete loss of DNA replication between 15 and 60 minutes post infection (mpi) (Figure 1F). These data are consistent with the observed loss of DNA replication upon antisense oligonucleotide (ASO) mediated depletion of Rhc in PhiKZ [15].

Expression of wild-type Rhc* largely complemented both phage titer and DNA replication (Figure 1E, F). Expression of the catalytic-dead D119N mutant partially rescued both phage titer and DNA replication (Figure 1E, F), in contrast to prior data showing that the PhiKZ Rhc catalytic-dead mutant (D102N) fully rescued phage titer and DNA replication [15]. Finally, we found that expression of the isolated Rhc RNase H or clamp domains did not rescue phage titer or DNA replication (Figure 1E, F). Overall, these data support a model in which Rhc contributes to DNA replication and the phage life cycle both through its RNase H catalytic activity, and through a second, non-catalytic role.

### The phage nucleus forms but doesn’t mature

To determine the stage of the replication cycle at which Rhc-knockdown infections are arrested, we first quantified the DAPI signal per nucleus as a proxy for nucleus size (Figure S3C). We found that the nucleus remained small compared to controls at 60 and 90 mpi, of comparable size to virions captured in the same data, which contain a single genome. From these data, we could not discern between these puncta representing the genome in the EPI vesicle or in the nucleus.

Next, we performed *in situ* cryo-electron tomography (cryo-ET) of Goslar-infected *E. coli* cells following Rhc knockdown (Figure 2). We acquired 30 tilt series from 17 cryoFIB-milled lamellae, revealing small, irregularly shaped, DNA-filled compartments that were physically separated from the host cytoplasm (Figures 2A,B; S5, S6). To quantify the volumes of these compartments within the boundaries of the lamellae, we developed a Continuity-Constrained, Slice-wise Volume Closure (CCSVC) algorithm. The algorithm combines geometric closure of segmentation gaps, cross-sectional continuity along the *z*-axis, and extrapolation of terminal regions to reconstruct three-dimensional volumes (Figure 2C; S7). At 60 mpi, these compartments had a median volume of approximately 0.013 µm^3^. Their distinctive single-layered, moniliform boundary with a periodically indented, beaded appearance was consistent with their identity as nascent phage nuclei rather than EPI vesicles. To further test this assignment, we performed subtomogram analysis of the boundary, yielding a cryoEM map at up to 22 Å resolution that revealed the assembly of chimallin tetramers forming the flexible proteinaceous shell of the phage nucleus (Figure 2D; S5). These observations indicate that Rhc knockdown arrests infection after phage DNA has been transferred into the nascent phage nucleus but before substantial DNA replication occurs. Together, our findings establish Rhc as an essential nuclear DNA replication factor in chimalliviruses and reveal that it contributes to DNA replication and phage development through both catalytic and non-catalytic functions.

**Figure 2:**
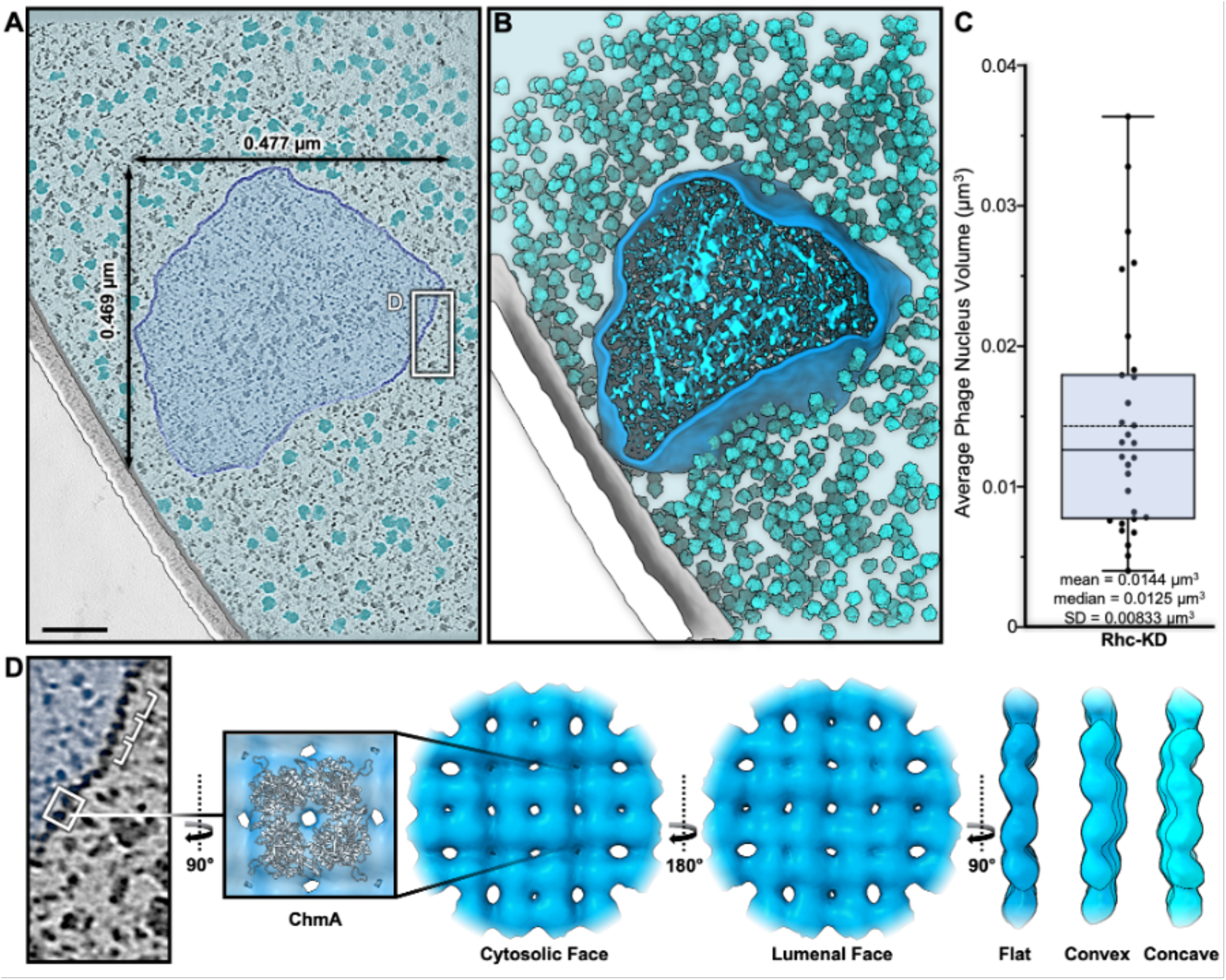
*In situ* cryoET of the Rhc-KD phage nucleus. **(A)** Tomographic slice showing the phage nucleus in an Rhc-KD *E. coli* cell 60 minutes after infection with Goslar. The phage nucleus is outlined and shaded in blue, with dimensions and are labeled. Ribosomes are shaded in cyan. Scale bar = 0.1 µm. See **Figure S6** for additional examples. **(B)** Segmentation of the tomogram shown in panel (A), showing the 3D volume of the bacterial cell (outer membrane white, inner membrane grey) with the phage nucleus (blue) surrounded by ribosomes (cyan). **(C)** Phage nuclei enclosed volume measurements over 30 nuclei using a Continuity-Constrained, Slice-wise Volume Closure (CCSVC) algorithm. Volume measurements show a mean of 0.0144 µm^3^ (dotted line) and a median of 0.0125 µm^3^ (solid line). **(D)** Subtomogram analysis of chmA forming the proteinaceous shell of the phage nucleus as seen in a tomographic slice of panel A. The atomic model of the Goslar chmA tetramer (pdb: 7SQU) is fitted into the cryoEM density. The chmA tetramers can arrange in a flat, convex or concave shell surface.

### Rhc is a DNA polymerase subunit

To gain insight into Rhc’s role in DNA replication, we sought to identify direct binding partners of the protein. First, we used an *in silico* protein-protein interaction screen based on AlphaFold 2 [17]. AlphaFold 2 produces low confidence models for most Goslar proteins, so we instead used proteins from PhiKZ to take advantage of more robust multiple sequence alignments for proteins from this phage, compared to Goslar. We predicted structures of 1:1 complexes of PhiKZ Rhc (gp155) with all other PhiKZ proteins, and identified three potential binding partners: DNAP^C^ (gp50), a predicted RuvC-like endonuclease (gp140), and the recently-identified phage nuclear import factor PicA [11,12] (gp69, also called Imp1; Figure 3B).

**Figure 3.**
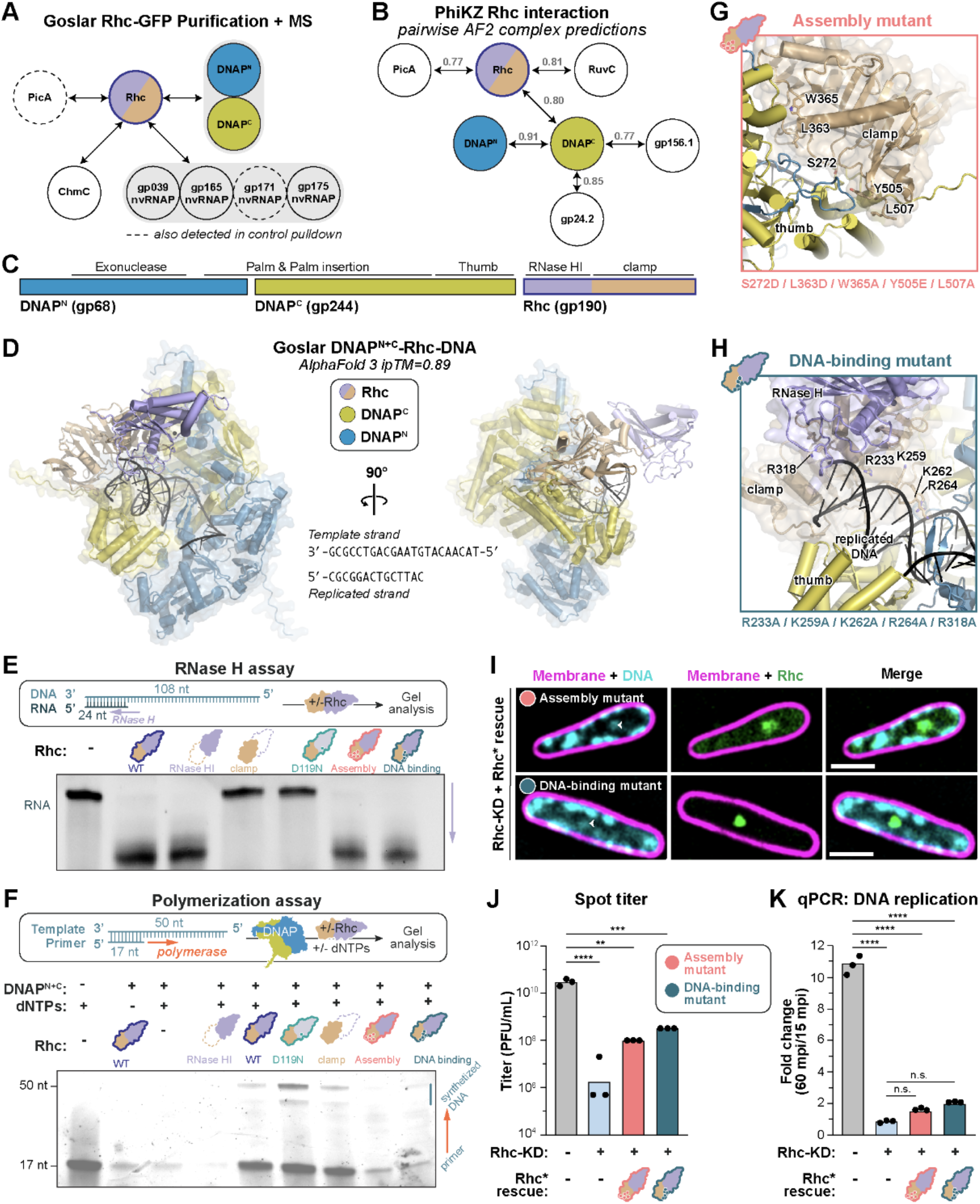
Rhc-DNAP/DNA interactions are required for DNA replication, and productive infection. **(A)** Proteins identified by mass spectrometry after purification of GFP-tagged Goslar Rhc from Goslar-infected *E. coli* cells (see **Table S2** for details). See **Figure S9A** for affinity pulldowns with purified Goslar proteins. **(B)** AlphaFold 2 interaction screen results with PhiKZ Rhc (gp155), DNAP^N^ (gp082), and DNAP^C^ (gp050). AlphaFold 2 interface predicted template modeling (ipTM) scores are shown for each protein pair; only the highest-ipTM partners for each protein are shown. **(C)** Domain organization of the Goslar DNA polymerase complex, showing DNAP^N^ (gp68), DNAP^C^ (gp244), and the RNase HI-containing Rhc protein (gp190). The predicted complex includes DNAP subunits including the exonuclease, palm and palm insertion, thumb, RNase HI, and clamp regions. **(D)** AlphaFold3 model of the Goslar DNAP^N+C^-Rhc-DNA complex (ipTM = 0.89), shown in two orientations. DNAP^N^, DNAP^C^, and Rhc are indicated, together with the modeled template and replicated DNA strands. **(E)** Purified WT Rhc, RNase HI, clamp,the catalytic-site mutant D119N, DNA binding and assembly mutants were purified and tested for RNase H activity using an RNA-DNA hybrid substrate. Reaction products were resolved by UREA gel electrophoresis to assess substrate cleavage. See **Figure S8A** for purification gels. **(F)** Primer extension assay with Goslar DNA polymerase complexes and the indicated DNA substrate. DNAP^N+C^ indicates the two-subunit gp244-gp068 complex (coexpressed and purified), and Rhc-WT, NTD, CTD, DNA-binding mutant, assembly mutant and D119N (catalytic dead mutant, separately expressed and purified, then mixed with DNAP^N+C^). See **Figures S8A, B** for purification gels. **(G)** Close-up view of the Rhc-DNAP^C^ interface. Residues predicted to mediate interaction between the Rhc C-terminal domain and the DNAP^C^ thumb domain (S272, L363, W365, Y505, L507) are shown. These residues were mutated (S272D, L363D, W365A, Y505E, L507A) to generate a DNAP-binding mutant (Assembly mutant). **(H)** Close-up view of the composite DNA binding surface formed by Rhc and the polymerase clamp and thumb domains. Basic residues within Rhc predicted to contact the replicated DNA duplex (R233, K259, K262, R264, R318) are highlighted. These residues were mutated to alanine (R233A, K259A, K262A, R264A, R318A) to generate a DNA-binding mutant. **(I)** Fluorescence microscopy analysis of Rhc-DNA and DNAP interaction mutants. Rhc knockdown cells complemented with Rhc mutants defective in DNAP interaction (Assembly mutant, **Figure 3G**) and DNA interaction (**Figure 3H**). Loss of Rhc-DNAP and DNA interactions disrupts nascent nucleus maturation. **(J)** Spot titer assay measuring infectious phage production under the indicated knockdown or complementation conditions. Titers are plotted as log_10_ PFU/mL. Individual biological replicates are shown as black dots. Statistical significance was assessed relative to NT control; n.s., not significant; **P < 0.01; ***P < 0.001; ****P < 0.0001. **(K)** Quantitative PCR analysis of phage DNA replication in Rhc-DNAP and DNA interaction mutants, measured at indicated time points and normalized first by 0 mpi and then to 15 mpi. NT control infections show robust genome amplification, whereas Rhc knockdown (Rhc KD) infections fail to support DNA replication. Disruption of the Rhc-DNAP interaction Rhc-DNA interaction does not restore genome amplification and remains indistinguishable from Rhc KD, indicating that both interactions are required for productive DNA replication. Bars represent mean ± s.e.m. (n = 3 biological replicates). Statistical significance was assessed by one-way ANOVA followed by Šídák’s multiple comparisons test (α = 0.05). Comparisons of NT versus all other conditions were highly significant (****P < 0.0001), whereas no significant differences were observed between Rhc KD and interaction mutants (n.s.). Scale bars: (I) 2μm.

Then, we expressed GFP-tagged Rhc in Goslar-infected *E. coli* cells, purified GFP-Rhc and associated proteins using anti-GFP affinity resin, then analyzed these proteins by mass spectrometry. Among the phage proteins we specifically detected in pulldowns of GFP-Rhc but not GFP alone were three subunits of the predicted non-virion RNA polymerase (gp039, gp165, and gp175; a fourth subunit gp171 was detected in both GFP-Rhc and GFP control pulldowns), along with the phage mRNA binding protein ChmC (gp176; Figure 3A, Table S2) [10]. We also detected two proteins with homology to DNA polymerases: gp68 (DNAP^N^) and gp244 (DNAP^C^) [13]. Overall, these data suggest that Rhc may associate directly with the phage RNAP and/or DNAP complexes. Supporting the validity of these data, a direct interaction between PhiKZ Rhc and PicA was recently reported [12]. Moreover, our pulldowns of Goslar GFP-Rhc also identified DNAP^C^ and PicA (gp244 and gp174, respectively; Figure 3A, Table S2, although PicA was also detected in the GFP-only control). Together, these data suggest that Rhc associates with the nuclear import factor PicA, the phage-encoded DNAP, and potentially a RuvC-like endonuclease.

To confirm these data and map the potential physical interaction between Rhc and the putative DNAP subunits, we separately expressed and purified His_6_-Rhc (full length, isolated RNase H domain, or isolated clamp domain, Figure S8) and GFP-tagged DNAP^N^ or DNAP^C^. In a GFP-affinity pulldown assay, we observed an interaction between DNAP^C^ and either full-length Rhc or its isolated clamp domain (Figure S9A). These data suggest that the Rhc clamp domain binds directly to the DNAP^C^ subunit of the phage DNA polymerase.

### Rhc boosts DNA polymerase activity

A concurrent study shows that PhiKZ Rhc binds a complex of DNAP^N^ and DNAP^C^, binds the replicated double-stranded DNA via its clamp domain, and boosts DNA polymerase activity of the complex *in vitro* (Basu *et al*., *bioRxiv*). An AlphaFold 3 [18]-predicted structure of DNA-bound Goslar DNAP^N^, DNAP^C^, and Rhc (Figures 3D, S10A) largely agrees with these data and highlights opposing surfaces on the Rhc clamp domain predicted to bind either DNAP^C^ or DNA (Figures 3G,H, S9B,C). To test whether Rhc contributes directly to DNA polymerase activity, we developed an *in vitro* polymerase assay using a 17-nt labeled primer and a 50-nt template strand.

First, we confirmed the RNase H activity of Rhc (Figure 3E) and then examined its effect on the DNAP^N+C^ complex. Addition of Rhc strongly stimulated DNA synthesis: whereas the DNAP^N+C^ complex alone exhibited prominent exonuclease activity, addition of wild-type Rhc shifted the reaction toward robust primer extension (Figure 3F), similar to the stimulatory effect reported for PhiKZ Rhc (Basu *et al*., *bioRxiv*). We next tested whether this stimulatory activity required the different functional features of Rhc. In addition to the catalytic dead mutant, we used the AlphaFold model to design and purify two mutants of Goslar Rhc predicted to disrupt binding to DNAP^N+C^ (S272D / L363D / W365A / Y505E / L507A; termed “assembly mutant”) or disrupt binding to DNA (R233A / K259A / K262A / R264A / R318A; termed “DNA-binding mutant”) (Figures 3G,H, S8A). Wild-type Rhc, the catalytic-dead D119N mutant, and the isolated clamp domain retained the ability to stimulate polymerization, whereas the RNase HI domain, the Rhc DNA-binding mutant, and the Rhc assembly mutant failed to enhance DNA synthesis (Figure 3F). These results indicate that stimulation of the DNAP complex does not require Rhc RNase H catalytic activity per se, but instead depends on features of the Rhc clamp domain that support DNA binding and association with the polymerase complex.

This finding matches our observation with genetics followed by light microscopy, qPCR and Spot titer assays (Figure 3I, J, K). The loss of DNA replication in the Rhc knockdown is partially complemented by expression of the catalytic-dead D119N mutant (Figure 1F). In the AlphaFold model, the RNase HI domain of Rhc is poised above the template DNA strand, such that its active site would be properly positioned to degrade an RNA strand annealed to the incoming template strand prior to replication. This positioning suggests that, in addition to providing a large DNA binding surface through its clamp domain, Rhc could degrade an RNA strand annealed to the template DNA to promote replication. This idea is consistent with our finding that the catalytic-dead D119N mutant does not fully complement DNA replication in Rhc-knockdown infections (Figure 1F).

To confirm DNA-binding and assembly mutation effects *in vivo*, we expressed the assembly and DNA-binding mutants in Rhc-knockdown Goslar infections. We observed that both mutants supported the formation of phage nuclei albeit generally much smaller than when complementing with wild-type Rhc and that both mutant proteins also localized to these small phage nuclei (Figure 3I). Expression of either mutant Rhc also partially rescued Goslar phage titer (Figure 3J, S10B), but did not significantly rescue DNA replication (Figure 3K).

Together, these data suggest that disrupting Rhc’s assembly into the DNA polymerase complex, or disrupting its ability to bind DNA within that complex, significantly hampers its ability to support DNA replication and phage development.

### Rhc regulates transcription independently of DNA replication

A major role of RNase H proteins across kingdoms is to support transcription by degrading R-loops formed when nascent RNA transcripts re-hybridize with the template DNA strand behind the RNA polymerase, displacing the non-template DNA strand [19–21]. To determine whether Rhc contributes to transcription, we performed RNA-seq in Rhc-knockdown Goslar infections and quantified genome-wide transcription relative to control (Figures 4A,B, S11A, Tables S5, S6, S8). Knockdown of ChmA prevents formation of the phage nucleus, resulting in the exclusive transcription of early genes from the EPI vesicle [7], and thus serves as a reference for the complete loss of nuclear transcription.

**Figure 4.**
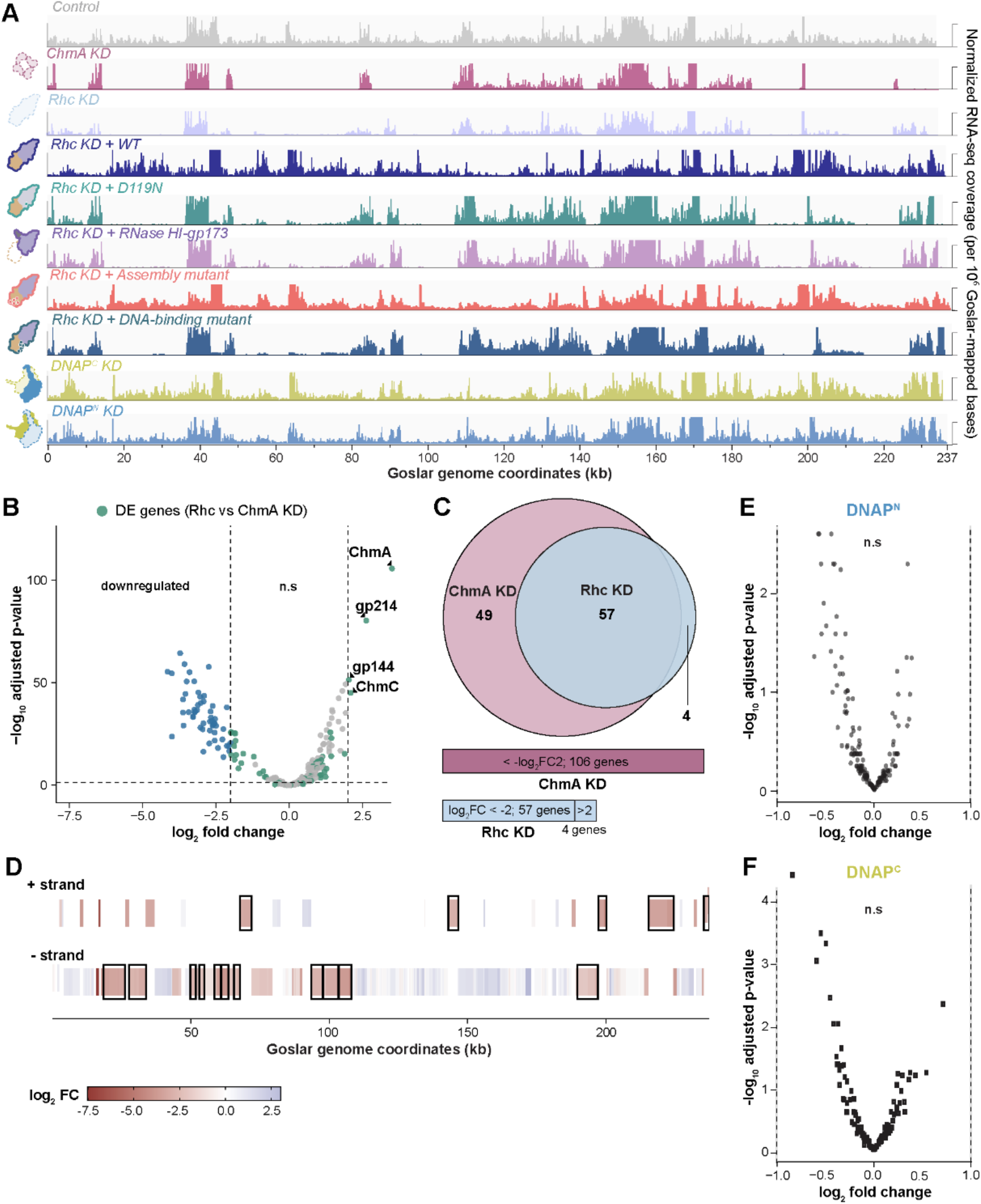
Rhc depletion broadly alters the phage transcriptional program independently of DNA polymerase abundance. **(A)** Genome-wide RNA-seq coverage across the 237-kb Goslar genome under the indicated infection conditions. RNA-seq reads were aligned to a combined *E. coli*-Goslar reference, and primary alignments with MAPQ ≥10 (mapping quality score) were retained for coverage analysis. Reads mapping to the Goslar genome were extracted, and coverage was normalized for each sample by the total number of Goslar-mapped bases using a scale factor of 10^6^/(total Goslar-mapped bases), thereby accounting for differences in sequencing depth and read length across datasets. Coverage was calculated in 50-bp bins, normalized by the total number of Goslar-mapped bases using a scaling factor of 10^6^/(total Goslar-mapped bases), exported as BigWig (.bw) files visualized in IGV at a fixed track height of 15. Three biological replicates are shown for each condition (**Figure S12A**), including the non-targeting control (Ctl.), ChmA knockdown, Rhc knockdown, complementation with WT, the catalytically inactive D119N mutant, gp173-tagged RNase HI, the DNA-binding mutant, assembly mutant, and depletion of the C- and N-terminal DNAP subunits (DNAP^C^ and DNAP^N^, respectively). ChmA knockdown RNA-seq data from our previous study [7] were reanalyzed using the same RNA-seq processing, alignment, filtering, and normalization pipeline applied to all other datasets in this study. **(B)** Volcano plot showing differential phage gene expression following Rhc depletion relative to the non-targeting control. Genes are plotted according to log2 fold change (x-axis) and −log10 adjusted P value (y-axis). Dashed lines indicate significance thresholds of |log2 fold change| ≥ 2 and adjusted P value < 0.05. Genes whose expression differs between Rhc KD and ChmA KD are highlighted in green. Selected genes showing increased expression following Rhc depletion are labeled. n.s, not significant. **(C)** Venn diagram showing the overlap between significantly differentially expressed genes identified following ChmA depletion and Rhc depletion. Genes were classified as differentially expressed using an adjusted P value less than 0.05 and an absolute log2 fold change greater than or equal to 2. 106 genes were down regulated in ChmA KD among 57 showed depletion in Rhc KD as well. 4 genes which were up-regulated in Rhc KD (Also see **Table S7** for details). **(D)** Gene-level log_2_ fold changes from Rhc KD relative to control are plotted across the Goslar genome, with genes separated by transcriptional strand. Contiguous same-strand groups of significantly downregulated genes are outlined to highlight Rhc-dependent transcriptional blocks. Rhc-dependent genes occur in discrete genomic regions rather than being uniformly distributed across the genome, consistent with coordinated effects on neighboring transcriptional units (Also see **Table S9** for details). Downregulated genes were defined as adjusted *P* < 0.05 and log_2_FC ≤ −2. **(E)** Volcano plot showing differential phage gene expression following depletion of DNAP^N^, relative to the non-targeting control. Dashed vertical lines indicate log2 fold-change cutoffs of −1 and +1. Neither perturbation produced a significant genome-wide transcriptional response at the indicated statistical thresh-old. n.s, not significant (see **Table S5**). **(F)** Volcano plot showing differential phage gene expression following depletion of DNAP^C^, relative to the non-targeting control (see **Table S5**).

In contrast to ChmA knockdown, in Rhc-knockdown infections we observed a partial effect on transcription, with 57 of 249 Goslar genes significantly downregulated, and four significantly up-regulated (adjusted p < 0.05, |log_2_FC| ≥ 2) (Figures 4A,B, S11A, S12, Tables S5, S7, S8). Most down-regulated genes in Rhc-knockdown infections are among the 106 late genes whose transcription is lost in ChmA knock-downs (Figure 4A,C). These data show that knockdown of Rhc specifically affects transcription of a subset of late genes that are transcribed in the phage nucleus by nvRNAP. Notably, the Rhc-dependent transcriptional defect was not randomly distributed across the genome. Downregulated genes frequently occurred in contiguous genomic blocks, and neighboring genes within inferred same-strand transcriptional units showed highly concordant changes in expression (Figure 4D, S13, Table S9).

In addition to the widespread loss of late gene transcription, Rhc depletion led to selective upregulation of a small subset of early genes, suggesting disruption of the normal temporal transcription program. Specifically, two early characterised genes, ChmA and ChmC, and two uncharacterised genes, gp214 and gp144, showed increased expression upon Rhc depletion (log_2_FC = 2.95, 2.10, 2.63, and 2.04, respectively).

Complementation of the Rhc knockdown with full-length Rhc* largely restored the wild-type transcriptional program. A small group of genes remained strongly reduced relative to control (log_2_FC ≤ −2), whereas a distinct subset showed localized overcompensation (log_2_FC ≥ 2) (Figures 4A, S11A,B,12, Table S10). In contrast, complementation with a catalytic-dead Rhc mutant (D119N) did not rescue transcription (Figures 4A, S11A,S12). Thus, in contrast to Rhc’s role in DNA replication, Rhc’s contribution to transcription depends on its RNase H catalytic activity. To test whether the RNase H domain is sufficient to support transcription, and since the isolated domain does not localize to the nucleus (Figures 1D, S4), we forced nuclear localization of this domain by fusing it to a known nuclear-localized protein, inactive gp173 deletant (Figure S14A). This construct localized to the phage nucleus (Figure S14B) but failed to rescue transcription and phage lysis (Figure 4A, S12, S14C), suggesting that Rhc’s clamp domain may also contribute to transcription. Supporting this finding, expression of the Rhc DNA-binding mutant also failed to rescue transcription, suggesting that the clamp domain’s ability to bind nucleic acids is important for Rhc’s role in transcription (Figure 4A, S12). Together, these experiments show that Rhc requires an active RNase H domain and nucleic-acid binding through its clamp domain, and that nuclear localization of the RNase H domain alone is not sufficient.

One model for Rhc knockdown’s effect on transcription is that transcription is indirectly affected by loss of DNA replication. To test this idea, we used CRISPRi-ART to knock down DNAP^N^ and DNAP^C^. Both knockdowns showed a loss of DNA replication as measured by qPCR (Figure S15A), but no Goslar gene met our significance criteria (adjusted P < 0.05 and |log_2_FC| ≥ 2) (Figures 4A,E,F, S12, S15B). In addition, the Rhc assembly mutant, which is impaired in its interaction with the DNA polymerase complex, successfully rescued the transcriptional phenotype (Figures 4A, S12), suggesting that full Rhc-DNAP interaction is not required for Rhc-dependent transcription. Together, these data separate the replication and transcription functions of Rhc. The DNA-binding and assembly mutants both support only small nuclei and fail to restore DNA replication (Figure 3I,K), yet only the assembly mutant restores transcription (Figure 4A, S12), so the transcriptional defect is not a secondary consequence of replication arrest or of failed nuclear growth. Rhc therefore supports transcription within the phage nucleus through a function that is separable from DNA replication and requires both RNase H catalytic activity and DNA engagement by its clamp domain.

## DISCUSSION

DNA replication in nucleus-forming chimalliviruses presents an unusual mechanistic problem. They encode two proteins with homology to DNA polymerases plus a DnaB-like replicative helicase, but lack a recognizable sliding clamp and primase proteins that are central to canonical bacterial and eukaryotic replisomes. Here, we show that Rhc resolves this problem by functioning as a clamp in addition to an RNase H by forming a three-protein assembly: the DNAP^N^ and DNAP^C^ proteins form a stable, extensively interdigitated heterodimer with overall architectural and catalytic similarity to B-family polymerases (Basu *et al*., *bioRxiv*), and a multidomain RNase H protein (Rhc) transiently associates with this enzymatic core and promotes DNA replication through both catalytic and non-catalytic means.

Notably, the Rhc-dependent genes include most of the annotated virion structure and assembly genes, whereas genes encoding the replication and repair machinery, including DNAP^N^, DNAP^C^, the DnaB-like helicase, a DNA helicase, DNA ligase, and RuvC remain expressed upon Rhc depletion. Thus, the complete loss of genome replication in Rhc-depleted infections cannot be readily explained by a secondary failure to express the replication machinery. Instead, this separation of phenotypes supports a direct role for Rhc in DNA replication, consistent with its physical association with DNAP^N+C^ and its ability to stimulate DNA polymerization *in vitro*.

Rhc’s C-terminal domain is distantly related to DNA sliding clamp proteins. While Rhc lacks the structural elements for oligomerization into a DNA-encircling complex, it retains an extensive DNA binding interface that binds the replicated double-stranded DNA emerging from the DNA polymerase active site (Basu *et al*., bioRxiv). Rhc promotes DNA replication independently of its RNase H catalytic activity *in vitro* and, at least in part, in infected cells, suggesting that it acts by stabilizing the binding of the DNAP on DNA, potentially boosting DNA replication processivity and/or catalytic rate. Consistent with this, mutants designed to disrupt the Rhc–DNAP interface or the Rhc–DNA interface both failed to stimulate DNA polymerization *in vitro*, and did not fully restore DNA replication *in vivo*. Both mutants nonetheless supported the formation of small phage nuclei and partially rescued phage titer; this may reflect incomplete disruption of the targeted DNAP- and DNA-binding interfaces.

Our data show that Rhc also contributes to DNA replication through its catalytic activity: unlike in PhiKZ, where a catalytic-dead Rhc mutant fully complements an ASO knockdown [15], we find that in Goslar the catalytic-dead Rhc only partially complements DNA replication and phage titer. Given the positioning of the Rhc RNase H domain adjacent to the DNA template strand in the three-protein polymerase complex, we speculate that Rhc may degrade RNA hybridized to the template DNA strand to promote replication.

Why do chimalliviruses possess an integrated RNase H in their DNA polymerases? Many organisms rely on RNase H proteins to degrade RNA primers during DNA replication, but these proteins are not integrated into the replisome. The best example of an integrated DNA polymerase-RNase H enzyme is in retroviral polymerases like HIV-1 reverse transcriptase: in this enzyme, a fused RNase H1 domain degrades the genomic RNA after reverse transcription into DNA, to allow second-strand DNA synthesis [22–25]. Chimalliviruses have a double-stranded DNA genome and therefore do not perform reverse transcription. We speculate that Rhc’s role may instead be the degradation of extended RNA-DNA hybrids that would otherwise stall DNA replication or cause replication-associated DNA damage, akin to the established mechanisms of R-loop mediated DNA break formation [26,27].

An examination of the chimallivirus life cycle provides one reason why R-loops might be particularly prevalent in these phages: the physical segregation of transcription from translation. In bacteria, a major mechanism for preventing R-loop formation is thought to be the tight coupling between transcription and translation, meaning that nascent transcripts are often actively translated (and thereby occupied by ribosomes) even while their transcription is progressing [28– 31]. In chimalliviruses, mRNAs are produced in the nucleus and must be transported to the cytoplasm for translation [1,9]. We have previously identified an mRNA-binding protein, ChmC, that is likely important for nuclear-cytoplasmic transport of mRNAs, and may also play a role in minimizing R-loop formation by binding mRNAs [10]. Nonetheless, the uncoupling of transcription from translation may leave chimalliviruses particularly prone to widespread R-loop formation, which could impact DNA replication and require an adaptation like Rhc to compensate. At the same time, the widespread presence of RNA-DNA hybrids in chimalliviruses could provide sites for the initiation of DNA replication, potentially explaining the lack of a dedicated primase in these phages.

Consistent with the idea that R-loops are widespread in chimalliviruses, we find that Rhc knockdown strongly affects the transcription of a large subset of late genes in Goslar. This role is independent of any effect on DNA replication, but requires both the RNase H catalytic activity and the nucleic acid binding interface on the protein’s clamp domain. Our data suggest that the clamp domain could help recruit Rhc to RNA-DNA hybrids in the absence of (or with reduced association with) the DNAP^N+C^ complex to degrade R loops and support transcription. The Rhc-dependent late genes cluster within long transcriptional units, supporting the possibility that transcript length contributes to their sensitivity to Rhc depletion. However, additional work will be required to determine whether transcript length itself drives this dependence, or whether other features such as transcriptional output, sequence composition, local genome topology, or propensity for R-loops formation are responsible.

Strikingly, we find that while Rhc is required for wildtype levels of late-gene transcription, loss of DNA replication does not reproduce the Rhc transcriptional phenotype. Knockdown of DNAP^N^ or DNAP^C^ results in no significant transcriptional changes, despite a complete loss of DNA replication. This counterintuitive finding suggests one of two possibilities: (1) transcription is limited to a single template genome in wild-type infections; or (2) the nvRNAP is the limiting factor in transcription, such that additional copies of the DNA template do not result in increased transcription. More work will be required to distinguish between these two possibilities, including analysis of any internal organization of the phage nucleus and possible sub-nuclear localization of the nvRNAP.

Fluorescence microscopy and qPCR show that Rhc knockdown produces small phage nuclei that contain a single genome copy, suggesting that growth of the phage nucleus is coupled to DNA replication, consistent with the growth of the nucleus alongside replication in wild-type infections [1]. This arrest is not obviously explained by a shortage of shell protein: ChmA transcript is elevated in Rhc-depleted infections, and Rhc knockdown in PhiKZ likewise raises ChmA protein levels [15]. The ChmA shell is a single-layer, flexible lattice that we had proposed grows by incorporation of soluble subunits into the existing lattice [8]. If incorporation continued without a matching increase in enclosed volume, the surplus shell would have to fold, giving a crumpled shell with a high ratio of surface area to DNA volume. Nuclear shells are irregular in shape even in wild-type infections [8], but we see no obvious signs of such excess folding in the small nuclei, which suggests that shell growth is limited by replication rather than by ChmA supply. Quantifying shell curvature in nascent and mature nuclei will allow this to be tested directly. Further, how the phage nucleus grows to accommodate DNA replication without allowing host-encoded immune nucleases to access the genomic DNA remains an open question. Whatever couples growth to replication, incorporation of new ChmA into the existing lattice could result in transient accessibility of the phage genomic DNA to host immune factors, potentially explaining the presence of DNA repair proteins and known anti-immune factors like AdfM in chimallivirus genomes [32].

We report the characterization of a dual-function RNase H enzyme in chimalliviruses, with independent roles in DNA replication and transcription (Figure 5). In DNA replication, Rhc stabilizes DNA polymerase on DNA and likely degrades RNAs annealed to the template strand, thereby promoting DNA replication in two ways. Rhc is also required for transcription of a subset of late genes, highlighting a likely role in R-loop degradation to support transcription of these genes. This work illustrates how large bacteriophages repurpose conserved enzymatic activities to overcome the unique challenges imposed by subcellular compartmentalization. Overall, by expanding the functional repertoire of RNase H, chimalliviruses have evolved a streamlined strategy to coordinate genome expression and replication within a nucleus structure, providing new insights into the minimal requirements for genome management and the evolution of complex viral replication strategies.

**Figure 5.**
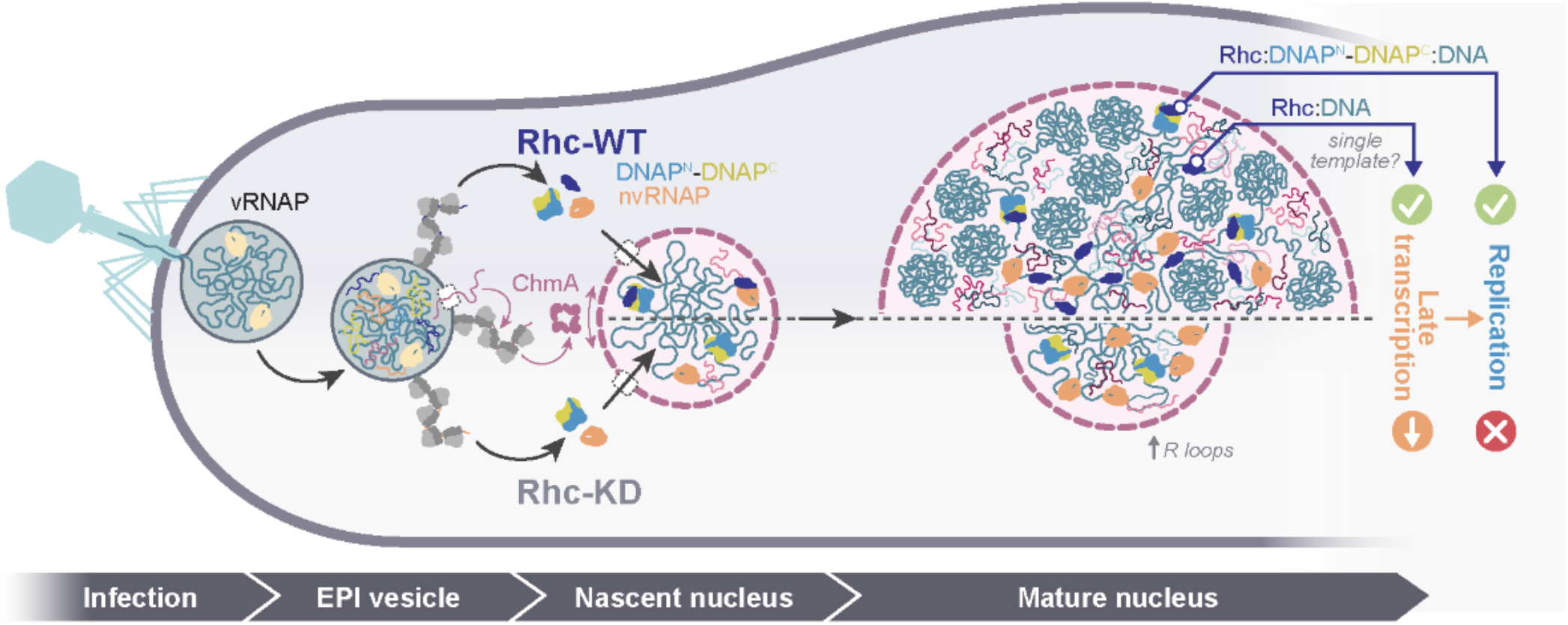
Model for Rhc-dependent genome replication, transcription, and phage nucleus maturation. Following infection, the virion RNA polymerase initiates early transcription within the membrane-bound early phage infection (EPI) vesicle. ChmA monomers subsequently assemble around the phage genome to form a nascent phage nucleus, while the non-virion RNA polymerase (nvRNAP) supports transcription of later phage genes. Rhc is recruited to the nascent nucleus, where it associates with the DNAP^N+C^ complex and promotes productive genome replication. Rhc activity also supports the late genes transcriptional program required for continued nuclear growth. Depletion of Rhc blocks DNA replication, alters phage transcription, and prevents expansion of the nascent compartment into a mature phage nucleus.

## MATERIALS AND METHODS

### Bacterial culture conditions and infection

The *E. coli* MC1000 cells were cultured in LB supplemented with appropriate antibiotics and incubated at 37 °C in a shaker incubator at 225 rpm. Then 100 µl of 0.3 to 0.5 O.D_600_ cells were plated on 25% LB, 1% agarose in a humidifier chamber and incubated at 37°C for 2-3 hr. The cells were infected with 100 µl of high titer Goslar lysate (plaque forming units, PFU/mL ≥ 10^10^; originally obtained from Johannes Wittman at the DSMZ) for specified time points. The infected and uninfected cells were collected and processed further for RNA or DNA isolation or making lysates for Western blotting.

### Goslar lysate production and plaque assays

To prepare Goslar lysates for infections, 0.5 mL of overnight wild-type *E. coli* MC1000 culture, 100 µL of high-titer Goslar lysate and 4.5 mL LB top agar (0.4%) were mixed and plated on 3-5 LB plates, then incubated overnight at 37°C. 5 mL of LB broth were added to each plate and allowed to sit for 4-5 hr at 30°C and all liquid was removed from the plate surface and centrifuged at 3000x g for 10 minutes. The supernatant was transferred to a new tube and cleared with chloroform (20 µL chloroform/5 mL lysate). The aqueous phase of the lysate was transferred to a fresh tube after incubating at room temperature for 10 minutes with periodic mixing by inversion. Lysate titers were 10^10^-10^11^ PFU/mL as measured by plaque assay on *E. coli* MC1000.

Plaque assays were performed in biological triplicate. 500 µL of overnight *E. coli* culture (MC1000 or MC1000 transformed with CRISPRi-ART knockdown plasmids) was mixed with 9.5 mL of 0.4% LB top agar containing antibiotics and inducing agents aTc (anhydrotetracycline; 100 nM) and IPTG (100 µM) and poured onto 20 mL LB plates containing the required antibiotics. Since the plates did not contain inducing agents, aTc was added to the top agar at initial concentrations to account for diffusion into the total final plate volume. Plates were spotted with a 10-fold dilution series of initially ~200 fold diluted Goslar from 10^1^-10^8^ (prepared in LB) in technical triplicate with 2 µL/spot and were dried. Once the spots dried, the plates were incubated at 37°C overnight. Plates were photographed on a white background using an iPhone16 Pro Max. Then, images were converted to 8-bit grayscale, and uneven illumination was corrected by generating a Gaussian-blurred duplicate of each image and dividing the original image by this background image. Contrast was then normalized using the same saturated-pixel setting for all images, followed by mild unsharp-mask sharpening. All images within a comparison group were processed using identical parameters, and no image-specific manual contrast adjustments were applied.

### Knockdown and complementation generation

The knockdown of the Goslar Rhc was generated and characterized as described earlier [7]. For knockdowns p15A-CmR vector, while for sfGFP complementation, vector pDSW206 [2]. were used respectively. All other plasmids were designed in Benchling and synthesized by GenScript.

We manually designed crRNA spacers, each 31 bp in length and reverse complementary to the 5’ end of the Goslar Rhc or DNAP^N/C^ ORFs, adhering to the design conventions as outlined earlier [16]. Goslar’s Rhc ORF ends 3 bp upstream of the gp191 gene, which has its own RBS (tggagt) 5 bp upstream of its ATG site indicating that these genes are translated independently (Figure S3). Our non-targeting control was a previously validated crRNA targeting the T4 phage major capsid protein [33]. Similarly, Goslar Rhc has its own RBS (atgagg) 6 bp upstream of the start codon (Figure S3). The lyophilized plasmids were diluted to 40 ng/µL and 1 µL diluted plasmid was added to electroporation competent MC1000 *E. coli*, which were prepared by washing in 10% glycerol, and electroporated with 1.6 kV using a BioRad electroporator. After transformants recovered in 1 mL LB broth at 37°C for 1 hr, they were spread on LB plates with appropriate antibiotics and incubated overnight at 37°C.

### Live-cell fluorescence microscopy

Imaging of uninfected and infected MC1000 cells was performed in biological triplicates. Host cells were grown to log phase (OD_600_ ~0.3-0.4) in LB broth with appropriate antibiotics. 10 µL were plated on single-well concavity slides containing 1% agarose and 25% LB imaging pads in single-well concavity slides. The pads contained appropriate antibiotics (100 µg/mL ampicillin, 30 µg/mL chloramphenicol, 100 nM aTc and 100 µM-2 mM IPTG to maintain the plasmid and induce expression of fluorescent protein fusions in complementations and 30 µg/mL chloramphenicol, 100 nM aTc in knockdowns). Cells were grown on the imaging pads for 2-2.5 hr at 37°C inside humidity chambers. For infection, 10 µL of Goslar lysate (PFU/mL ≥ 10^10^) were spread on the pad surface and the slide was incubated at 37°C within the humidity chamber until the desired timepoints.

Before imaging, pads were stained with 8 µL of 25 µg/mL DAPI (Thermo Fisher Scientific Cat# D1306) and 3.75 µg/mL FM4-64 (Thermo Fisher Scientific Cat# T13320) at room temperature. All microscopy was performed on a DeltaVision Elite Deconvolution microscope (Applied Precision, Issaquah, WA, USA) via DeltaVision SoftWoRx (version 6.5.2). Images were deconvolved in DeltaVision SoftWoRx (version 6.5.2) and analysed using Fiji.

DAPI signal per nucleus was quantified in Fiji (version 2.16.0/1.54p) from the DeltaVision images of live Goslar-infected *E. coli* MC1000 cells (non-targeting control, NT, and Rhc-KD) described above, acquired as z-stacks of nine slices at 0.2 µm spacing (15.6578 pixels/µm, 0.064 µm per pixel). In a first experiment, Rhc-KD and NT infections were imaged at 60 mpi (Rhc-KD and NT set 1); in a second experiment, NT infections were imaged at 60 mpi (set 2) and 90 mpi; the images shown in the figures are from 60 mpi. Free virions from the inoculum, located outside the cells in the same images, were measured as the single-genome reference and pooled across the Rhc-KD, NT 60 mpi and NT 90 mpi images (n = 31, 36 and 32, respectively; 99 in total). For each nucleus or virion, the central z-slice, defined as the middle of the DAPI-positive z-range, was selected, a region of interest (ROI) was outlined manually on the DAPI channel, and area (µm^2^) and mean gray value were measured with Analyze > Measure. Background was the mean gray value of cell-free pad, measured once per dataset (138.3, 147.6, 123.4 and 143.6 a.u. for Rhc-KD, NT 60 mpi set 1, NT 60 mpi set 2 and NT 90 mpi, respectively). Integrated DAPI intensity was calculated for each object as ROI area × (mean gray value − background) and divided by the median integrated intensity of the virions (virion equivalents). Data are from three biological replicates with n = 99 free virions, 72 Rhc-KD nuclei, and 133, 100 and 101 NT nuclei (NT 60 mpi sets 1 and 2 and NT 90 mpi, respectively). Because virion ROIs span only a few pixels and may capture only part of the point-spread-function signal, NT values in virion equivalents may be over-estimated by up to about two-fold; the Rhc-KD to virion ratio is less affected. Because the measurement uses a single z-slice, manually drawn ROIs, live staining without equilibration and datasets acquired in separate sessions, values are semi-quantitative and are compared across groups only as relative differences.

### Quantitative polymerase chain reaction (qPCR)

Cells were collected at 0, 15 and 60 mpi, pelleted, and washed with 1X PBS (Gibco, 10010072). The pellets were stored at −20°C after snap-freezing in liquid nitrogen. The pelleted cells were resuspended in 200 µL 1X TE (10 mM Tris-HCl pH 8.0, 1 mM EDTA pH 8.0) and mixed with 100 µg of lysozyme, followed by incubation at 37°C for 1 hr. DNA was isolated using a Quick-DNA Miniprep kit (Zymo Research, D4068). The concentration of genomic DNA was analyzed using a Quibit 4 Fluorometer (Invitrogen). For qPCR, the DNA was diluted to 10 ng/µL, and the qPCR reaction was set up using Luna Universal qPCR Master Mix (NEB, M3003). The data were normalized first by 0 mpi and then by 15 mpi where required. The bar graph was generated using GraphPad Prism (version 10.0.0; GraphPad Software) for three independent biological replicates. The significance was calculated using one way ANOVA tests with 95% confidence intervals. As negative control, the same experiment was performed in non targeting control. Primer sequences used for qPCR are listed in Table S4.

### GFP-trap purification and mass spectrometry

GFP or GFP-tagged Goslar Rhc was expressed in *E. coli* MC1000 from vector pDSW206, and cells were infected with Goslar, and plated on 1% agarose, 25% LB,100 µM IPTG plates. Cells were collected at 60 minute post infection (mpi) using 1 mL LB, pelleted, and washed with 1X DPBS (Gibco). The cell pellets were dissolved in 200 µL of lysis buffer (10 mM Tris-HCl pH 7.5, 150 mM NaCl, 0.5 mM EDTA, 0.5% NP-40 substitute, 0.09% sodium azide) and sonicated to lyse the cells. The cell debris was removed after spinning the cells at 13000x g for 30 min at 4°C. Then in the clear cell supernatant, 300 uL of dilution buffer (10 mM Tris-HCl pH 7.5, 150 mM NaCl, 0.5 mM EDTA, 0.018% sodium azide) and 25 µL of dilution buffer-equilibrated GFP-Trap Agarose beads (Chromtek, cat# gta) were added. The sample was incubated at 4°C for 1 hr on a rotator. The beads were collected by centrifugation at 2500x g for 2 min at 4°C. The collected were washed three times in a wash buffer (10 mM Tris-HCl pH 7.5, 150 mM NaCl, 0.05 % NP40 substitute, 0.5 mM EDTA, 0.018% sodium azide). The protein was eluted using 100 µL of acidic elution buffer (200 mM glycine pH 2.5) and immediately neutralized with 10 µL of 1 M Tris-HCl pH 10.4. Then samples were diluted in TNE (50 mM Tris-HCl pH 8.0, 100 mM NaCl, 1 mM EDTA) buffer. RapiGest SF reagent (Waters Corp.) was added to the mix to a final concentration of 0.1% and samples were boiled for 5 min. TCEP (Tris (2-carboxyethyl) phosphine) was added to 1 mM final concentration and the samples were incubated at 37°C for 30 min. Subsequently, the samples were carboxymethylated with 0.5 mg/mL of iodoacetamide for 30 min at 37°C followed by neutralization with 2 mM TCEP (final concentration). Proteins samples prepared as above were digested with trypsin (trypsin:protein ratio −1:50) overnight at 37°C. RapiGest was degraded and removed by treating the samples with 250 mM HCl at 37°C for 1 hr followed by centrifugation at 14000 rpm for 30 min at 4°C. The soluble fraction was then added to a new tube and the peptides were extracted and desalted using C18 desalting columns (Thermo Scientific, PI-87782). Peptides were quantified using BCA assay and a total of 1 µg of peptides were injected for LC-MS analysis. Trypsin-digested peptides were analyzed by ultra high pressure liquid chromatography (UPLC) coupled with tandem mass spectroscopy (LC-MS/MS) using nano-spray ionization. The nanospray ionization experiments were performed using a Tim-sTOF HT pro hybrid mass spectrometer (Bruker) interfaced with nano-scale reversed-phase UPLC (EVOSEP ONE). Evosep method 30 samples per day was utilized using a 15 cm × 150 µM reverse-phase column packed with 1.5 µM C18-beads (PepSep, Bruker) at 58°C. The analytical columns were connected with a fused silica ID emitter (10 µM ID; Bruker Daltonics) inside a nanoelectrospray ion source (Captive spray source; Bruker).For the timsTOF HT settings, the following parameters were adapted, starting from the PASEF method for standard proteomics. The values for mobility-dependent collision energy ramping were set to 95 contrast an inverse reduced mobility (1/*k*_0_) of 1.6 V s/cm^2^ and 23 eV at 0.73 V s/cm^2^. Collision energies were linearly interpolated between these two 1/*k*_0_ values and kept constant above or below. No merging of TIMS scans was performed. Target intensity per individual PASEF precursor was set to 20000. The scan range was set between 0.6 and 1.6 V s/cm^2^ with a ramp time of 166 ms. 14 PASEF MS/MS scans were triggered per cycle (2.57 s) with a maximum of seven precursors per mobilogram. Precursor ions in an *m*/*z* range between 100 and 1700 with charge states ≥3+ and ≤8+ were selected for fragmentation. Active exclusion was enabled for 0.4 min (mass width 0.015 Th, 1/*k*_0_ width 0.015 V s/cm^2^). Protein identification and label free quantification was carried out using Peaks Studio 12 (Bioinformatics solutions Inc.)

### AlphaFold 2 protein-protein interaction screening

Protein-protein interactions in the PhiKZ proteome were predicted using AlphaFold 2 coupled with AlphaPulldown [34]. All annotated PhiKZ protein sequences were retrieved from the NCBI reference database (NC_004629.1) and used to construct a comprehensive set of pairwise interaction predictions for all possible protein pairs. Multiple sequence alignments and structure predictions were generated using AlphaFold 2 multimer. Jobs were parallelized across GPUs on a high-performance computing cluster to enable large-scale screening of pairwise interactions. For each predicted complex, model confidence was assessed and passed through high-stringency filters (Predicted Aligned Error (PAE) < 7 Å, predicted TM-score (pTM) > 0.7, interfacial TM-score (ipTM) > 0.65).

### Protein expression and purification

To purify Goslar Rhc (gp190) and its different domains, we designed a His_6_-tagged construct in 2BT plasmid (UC Berkeley Macrolab vector 2-BT, Addgene #29666) using Benchling. The cloning was performed by Genscript. The plasmids were transformed into in-house prepared electrocompetent *E. coli* Rosetta2 pLysS cells. 4×500 mL cultures were grown in 1 L flasks at 37°C to an A_600_ of 0.3, induced with 0.3 mM IPTG at 16°C overnight. Cells were harvested by centrifugation at 4000 rpm at 4°C and resuspended in 20 mL of Buffer A (20 mM Tris-HCl pH 8.0, 100 mM NaCl, 5% glycerol, 10 mM imidazole, Protease inhibitor cocktail (Millipore sigma, cat# 11836170001). Then cells were sonicated and cell debris was removed at 10000x g for 30 min followed by centrifugation at 100,000x g for 2 hrs. Supernatant was collected and incubated with Ni-sepharose beads (Cytiva, cat # 17526801) overnight in 4°C on rotation. Next day, collected the flow-through by passing through manual gravity flow column (Cytiva, cat#17-0435-01). Ni-sepharose beads were washed with 50 mL of buffer B (20 mM Tris-HCl pH 8.0, 100 mM NaCl, 5% glycerol, 40 mM imidazole, Protease inhibitor cocktail (Milliporesigma, cat# 11836170001)). 30 fractions were collected with 15 mL of buffer C (20 mM Tris-HCl pH 8.0, 100 mM NaCl, 5% glycerol, 400 mM imidazole, Protease inhibitor cocktail (Milliporesigma, cat# 11836170001). The presence of protein in the fractions were analysed by SDS-PAGE analysis with Coomassie Brilliant Blue R-250 Staining Solution (BioRad, cat # 1610436). The fractions containing protein of interests were pooled together dialysed with buffer D (20 mM Tris-HCl pH 8.0, 100 mM NaCl, 5% glycerol, Protease inhibitor cocktail (Milliporesigma, cat# 11836170001) using Amicon ultra centrifugal filter (Milliporesigma, UFC903008). The purified and dialysed proteins were aliquoted and stored in −80°C. For the DNA-binding and assembly mutants, the same purification protocol described above was followed with minor modifications, including induction with 1 mM IPTG and an additional size-exclusion chromatography step. Following Ni-affinity purification, the eluted protein was concentrated and further purified on a Superdex 200 Increase 10/300 GL column (Cytiva) equilibrated in SEC buffer containing 20 mM Tris-HCl (pH 7.5), 250 mM NaCl, and 2 mM β-mercaptoethanol. Fractions containing the target protein were pooled, concentrated, and stored at −80°C.

Goslar DNAP subunits gp068 (DNAP^N^; see sequence below) and gp244 (DNAP^C^; NCBI accession YP_009820875.1) were cloned into the UC Berkeley Macrolab vector 2-BT (Addgene #29666). Constructs were designed in Benchling and synthesized by GenScript. The annotated gp068 coding sequence (NCBI accession YP_009820753.1) lacked the C-terminal region because the gene is interrupted after the predicted active site by a self-splicing intron containing the homing endonuclease gene gp069. To infer the full coding sequence, we used a group of homologous sequences (99.14% identical along their entire length) available in NCBI, exemplified by accession EKA1377486.1. The inferred full coding sequence for Goslar gp068 is bases 79690-81186 and 82949-83203 of NCBI RefSeq NC_048170.1, with the following protein sequence (residues not annotated in accession YP_009820753.1 denoted by an underline):

>gp068 MSEQERKPKKKLLGIQCRHISYNE-ANDGSKNDLMIAKEIM-FFDDNTQATNLRFFENYKRPFWVTVESKRKNKYKKESE KLSHVR-RYDVRQSELVEKISSVLYKRRADDNTRLQQLARSPYLYG TDVSASCYLKTQYNTKWPEYVDKLPASIACFD-IETDVVHGHGKIVCMSITMGYNAIL-CYTKDFVGHIPNAIEKTHEKIRYYLAERLNTKEDPHAFE KYHYDVRIVESPAQAVIQCMQKVHEWQPDYLLGWNI-AFDIPRMLQALNEE-GYRPEDVFSDPRVPPKYRIARWVPPKTDKVSASGRKTNL TLEQTWPMFYTMATYQVMDAMCGYKFVRLGRPDEGR YSLDAVLERNKCHTKLKAIPG-TEGFTELAWHEEMQTNHQIDY-GVYNLWDNQGIEALDQKTKDYSMTIPIRLAASDLKDFSS NPKRICDKLFFVYLNRGEVMGCTADDMTQEEDKWVIS RTGHIVTLPCDLMADNGIHLFWDMP-NLRSYIRLFVADLDISSS-YPNGQILLNLSRACVFRELSRIEGVSDYQFRKLGINFITGGKTNAVDFCRIVYKAPSMFELLEHFDATRPKGEARTIE

For expression of DNAP^N+C^, vectors encoding His_6_-DNAP^C^ and untagged DNAP^N^ (with ampR replaced with smR gene under same promoter) were co-transformed into *E. coli* Rosetta2 pLysS cells, then 1 L cultures in 2 L flasks were grown at 30°C to an A_600_ of 0.4, induced with 0.5 mM IPTG, then grown 16-18 hours at 16°C. Cells were harvested by centrifugation and resuspended in Buffer A (20 mM Tris-HCl pH 7.5, 300 mM NaCl, 10% glycerol, and 2 mM β-mercaptoethanol) plus 10 mM imidazole. Cells were lysed by sonication in the presence of protease inhibitor cocktails (Millipore sigma, cat# 11836170001). Cell debris was pelleted by centrifugation at 20000 g for 30 min at 4°C, and proteins were passed over a Ni^2+^ affinity chromatography column (Ni Sepharose, Cytiva 17-5318-02), washed with Buffer A plus 40 mM imidazole, and eluted with Buffer A plus 400 mM imidazole. Eluted protein was concentrated, dialyzed and stored in Buffer A containing 5% glycerol at −80°C.

For GFP trap, GFP-tagged Goslar DNAP constructs were expressed in *E. coli* MC1000 using the plasmid vector pDSW206. Cells were infected with Goslar phage and plated on 1% agarose, 25% LB plates supplemented with 0.1 mM IPTG to induce expression of the GFP-tagged proteins.

Infected cells were harvested at 60 mpi by resuspending the infected lawn in 1 mL LB medium. Cells were pelleted by centrifugation, washed once with 1× DPBS (Gibco). Cells were lysed by sonication in 200 µl of DNaseI buffer, and the cell debris was removed by using centrifugation at 13,000*x*g for at 4°C for 30 min. Then the lysate was resuspended was incubated with DNase I for 1 hr at 37 °C.300 µl lysis buffer containing 10 mM Tris-HCl (pH 7.5), 150 mM NaCl, 0.5 mM EDTA, 0.5% Nonidet™ P-40 substitute, and 0.09% sodium azide was added to the sample followed by 300 µl of dilution buffer(10 mM Tris-HCl pH 7.5, 150 mM NaCl, 0.5 mM EDTA, and 0.018% sodium azide). 2 µg of purified (Figure S8) His-tagged Rhc (active site mutant D119N) and CTD were subsequently added to the clarified lysates and incubated for 1 hr at 4°C with gentle rocking to allow complex formation. Then the samples were incubated with 25 µL GFP-Trap agarose beads (Chromotek) pre-equilibrated in a dilution buffer for 1 hr at 4°C with rotation. Beads were collected by magnetic rack at 4°C and washed three times with wash buffer containing 10 mM Tris-HCl (pH 7.5), 150 mM NaCl, 0.05% Nonidet™ P-40 substitute, 0.5 mM EDTA, and 0.018% sodium azide. Bound proteins were eluted using 100 µL acidic elution buffer (200 mM glycine, pH 2.5) and immediately neutralized with 10 µL 1 M Tris-HCl (pH 10.4). Eluted samples were probed for GFP and His tags using Jess Western blot as described Below.

### Western blot using Jess

Samples were resuspended in 2x SDS loading buffer (0.1 M Tris HCl, 0.004% bromophenol blue, 4% SDS, 20% glycerol, pH 6.8) and 5% BME. Then, the samples were boiled for 3-4 min at 95°C and vortexed for 30 seconds. Western blots were performed using the ProteinSimple Jess Automated Western Blot System according to its “multiplexing” immunoassay procedure. Samples were re-boiled, vortexed, and then diluted to an optimal concentration using a 0.1 dilution of 10X Sample Buffer (ProteinSimple) and antibodies were prepared by dilution with Antibody Diluent (ProteinSimple). Both samples and antibodies are diluted to the ideal concentration at which the targeted proteins stayed within linear dynamic range of the assay and minimized baseline signal noise. Phage-encoded protein targets were detected using an HRP-conjugated anti-Histidine tag (Biolegend #652504) and an anti-rabbit anti-GFP tag (Invitrogen #A10260). The secondary used to visualize bands for Goslar proteins with GFP tags was an anti-rabbit chemiluminescent antibody (Protein-Simple). The samples and antibodies were loaded into 12-230 kDa plates which are preloaded with a separation matrix, a stacking matrix, running buffer, and matrix removal buffer. Separation Modules with 8×25 Capillary Cartridges (ProteinSimple) were used for these assays into which the matrix, samples, and antibodies are aspirated. The Jess immunoassay protocol puts samples through separation, immobilization, immunoprobing, detection, and quantitation of chemiluminescence signals from each target protein.

### RNase H assays

RNase H activity was assayed using RNase H Reaction Buffer (New England Biolabs, NEB). The substrate was prepared by annealing a 108-nt DNA oligonucleotide with a 24-nt Cy5-labeled RNA oligonucleotide. The oligonucleotides were heated at 95°C for 10 min and then slowly cooled to allow duplex formation. The annealed substrate was used at a final concentration of 250 nM. RNase H reactions were incubated at 37°C for 30 min in a Bio-Rad PCR thermocycler. Reactions were stopped by addition of 2XTBE-Urea Sample Buffer (G-Biosciences, cat # 786-474) and heating at 95°C for 5 min. A total of 10 µL of each reaction was loaded onto a 15% Mini-PROTEAN TBE-Urea polyacrylamide gel (Bio-Rad, cat # 4566053). Gels were electrophoresed at 170 V for 40 min and subsequently imaged with the Cy5 channel on a Bio-Rad Chemidoc MP imager. DNA sequence used: 5′-CGAATCAGTGAATAACACCGCATT-GCCCGCTGCCTCCACAGCTTCGGGTGTTAACGAC-GCATTATTCTCCTGAATGGTTTGAATAGCTTGGTCGTAGTAATTAGACAT RNA sequence used: 5′(Cy5)-AUGUCUAAUUACUACGACCAAGCU

### In vitro polymerase assay

DNA polymerase activity assays were performed as previously described [35]. Briefly, a template DNA strand was annealed to a 5′-6-FAM labelled DNA primer (see below for sequences) by mixing equimolar quantities in annealing buffer (10 mM Tris, pH 7.5, 50mM NaCl, 1mM EDTA), heating to 95°C for 5 minutes, then slow-cooled over 30 minutes in a PCR machine. All reactions were performed with 25 nM annealed template-primer, 300 nM Goslar DNAP^N+C^ and or Rhc (WT, D119N, Clamp,RNase HI, DNA-binding mutant and assembly mutant), and 200 µM each dNTP in polymerase assay buffer (50 mM Tris, pH 7.5, 1 mM DTT, 2 mM MgCl_2_). 10 µL reactions were incubated at 25°C for 12 min, then reactions were stopped by the addition of 10 ul of TBE-Urea Sample Buffer (G Biosciences, cat# 786-474) and heated at 95°C for 5 min, followed by separation on15% Mini-PROTEAN TBE-Urea polyacrylamide gel (Bio-Rad, cat # 4566053). Gels were electrophoresed at 170 V for 40 min and subsequently imaged with the Cy2 channel (492 nm excitation) on a Bio-Rad Chemidoc MP imager. DNA sequences used: Template: 5′-CGGTACCCGGGGATCCTCTAGAGTCGAC-CTGCAGGCATGCAAGCTTGGCA-3′ Primer: 5′-(6-FAM)-TGCCAAGCTTGCATGCC-3′

### CryoET sample preparation and data acquisition

*E. coli* cells were grown in LB medium to an OD_600_ 0.4 and 100 µL were spread on 1% agarose, 25% LB, 6 cm diameter plates. After incubating at 37°C for 3 hr inside a humidor, cells were infected with Goslar lysate and incubated again at 37°C. At 60 mpi, cells were collected and centrifuged at 6,000 *x* g for 30 s. The cells were then resuspended in LB medium to an OD_600_ of 4. A 4 µL volume of cells was applied to R2/1 Cu 200 mesh Quantifoil grids that had been glow-discharged for 1 min at 0.19 mbar and 20 mA using a PELCO easiGlow device. After backside blotting for 6s, grids were plunge frozen in a 50:50 ethane:propane mixture using a custom-built manual plunging device (Max Planck Institute of Biochemistry). Samples were cryoFIB milled with an Aquilos 2 dual beam microscope (Thermo Fisher Scientific) at a 9° angle to generate ~150-200 nm thick lamellae. CryoET data was collected on a Titan Krios G3 (Thermo Fisher Scientific) operated at 300 keV equipped with a K3 detector and a 1067HD BioContinuum energy filter (Gatan) with 15 eV slit-width. Tilt series were acquired using the dose-symmetric scheme with a +/-54° tilt range and 3° increments at a pixel size of 1.695 Å and a 3-4 µm nominal defocus. Data was acquired automatically using SerialEM v4.2-beta [36] in combination with parallel cryo electron tomography (PACE-tomo) [37]. Tilt series were collected at 4.05 e^-^/Å^2^/tilt, resulting in a total fluence of 150 e^-^/Å^2^. Tomograms were reconstructed at a final pixel size of 6.78 Å using Warp v2.0.0dev31 [38] and Etomo v4.11.25 [39] patch tracking.

### CryoET data analysis and phage nuclei volume calibration

For representation, tomograms were deconvolved using Warp v2.0.0dev31 or denoised plus CTF and missing wedge corrected using IsoNet2 v2.0.0-beta [40]. For tomogram segmentation, membranes and phage nuclear densities were created using MemBrain v0.0.9 [41], Paintera v1.13.6 [42] and IsoNet2 v2.0.0-beta.

To reconstruct and measure the enclosed volume from phage nuclei or, more generally, segmented structures containing discontinuities and open boundaries, we developed a Continuity-Constrained, Slice-wise Volume Closure (CCSVC) algorithm. The method operates on binary segmentations independently in each tomographic z-slice while using cross-sectional continuity along the z-axis to select among alternative closures. 30 tomograms containing phage nuclei were first segmented with MemBrain v0.0.9 and manually cleaned up in UCSF ChimeraX v1.9 [43] before applying the CCSVC algorithm and measuring phage-nuclear volumes.

For each z-slice, the phage-nucleus segmentation was converted to a binary mask and skeletonized. Connected components of the skeleton were identified, and segmentation endpoints were defined as skeleton voxels having a single connected neighbor. Candidate closures were generated by pairing endpoints belonging to separate portions of the segmented nucleus. Endpoint pairs were connected by straight voxel paths, producing closed boundaries for subsequent filling. For each candidate closure, the enclosed cross-sectional region was determined by binary hole filling. The resulting mask was subsequently subjected to a one-voxel erosion followed by a second hole-filling operation. This procedure reduced the influence of artificial connecting paths while retaining the enclosed region defined by the candidate boundary. The number of voxels contained within the resulting cross-section was used as a measure of its cross-sectional area. Because multiple endpoint pairings can produce geometrically plausible closures within a single slice, candidate solutions were evaluated using information from neighboring z-slices.

For each candidate, the resulting cross-sectional area was compared with the areas of the nearest valid preceding and following slices. Solutions producing abrupt changes in cross-sectional area were penalized, whereas solutions yielding a smoother progression of area along the z-axis were favored. A secondary penalty was applied to the length of artificial connections, favoring shorter connections when alternative closures produced comparable cross-sectional continuity. Candidate selection was iteratively refined along the z-axis to reduce discontinuities in the reconstructed cross-sectional area profile. Where no valid closure could be obtained for an individual slice, the cross-sectional solution from the nearest reliable slice was propagated to maintain continuity of the reconstructed volume.

In addition to local discontinuities within the segmentations, the segmented phage nuclei may terminate at a boundary of the available tomographic volume due to FIB milling. Such terminal regions may represent open boundaries of the segmentation rather than a physical termination of the underlying structures. They were therefore treated separately from local slice-wise closure. For the leading terminal region, a reliable cross-sectional region within the segmented volume was selected as an anchor following the strongest initial increase in cross-sectional area. This anchor region was propagated toward the terminal boundary using the displacement of the segmentation centroid relative to the anchor slice. During propagation, the predicted cross-sectional region was constrained by the convex hull of the segmentation pixels in each target slice to maintain consistency with the observed geometry. The resulting regions were filled to produce a continuous terminal extension. This step represents a geometric extrapolation of the segmented volume beyond the reliably observed region, rather than a local closure of a segmentation gap.

The resulting closed cross-sectional masks were combined along the z-axis to generate a three-dimensional binary volume of the phage nuclei in each of the 30 tomograms. The reconstructed volume was calculated from the number of voxels contained within the final mask and the calibrated voxel dimensions:

V=N_vox_d_x_d_y_d_z_,

where N_vox_ is the number of voxels assigned to the reconstructed volume and d_x_, d_y_ and d_z_ are the voxel dimensions along the three axes.

### Subtomogram analysis

For representation and segmentation, 4,505 ribosomes were picked with crYOLO v1.9.4 [44] using a pretrained model. Ribosomes were extracted at a pixel size of 6.78 Å and 3D classified with Relion v3.1.4 [45] to clean up the dataset. Clean particles were then re-extracted at a pixel size of 3.39 Å and 3D refined to a resolution of 17 Å with further subsequent 3D refinement applying a mask to the entire ribosome volume. The resulting cryoEM map at a resolution 13 Å was used as input for further refinement with Warp/M, leading to a final cryoEM map for the overall ribosome at 9.3 Å. The FSC curve shows oscillations, most likely due to particles originating from too few tilt series that do only provide similar nominal defoci. For representation, the map was then projected back into the tomograms with their coordinates and angular orientations using the UCSF ChimeraX v1.9 plugin ArtiaX v0.5.0 [46].

To refine a ChmA cryoEM map of the proteinaceous phage nuclear shell, 1,828 particles were picked using GAP-STOP v0.4 [47] based on the STOPGAP [48] algorithm using emd25395 as a template. Particles were extracted at a pixel size of 6.78 Å and pre-oriented perpendicular to the surface of the nuclear shell using a dmodels.vesicle [49] model. The first 3D refinement with angular sampling restriction and C4 symmetry in Relion v3.1.4 resulted in a resolution of 30 Å. Subsequent 3D classification without image alignment and a T=1 value resulted in 3 classes. The particles in one class were recentered onto the chmA tetramer and again 3D refined with a C4 symmetry, this time not restricting angular sampling to give more freedom for alignment. Particles of all classes were now combined and manually cleaned up by removing duplicates and by reorienting particles to be perpendicular to the nuclear shell. The clean dataset of 886 particles was again 3D classified. This resulted in 3 classes representing chmA arranged in a flat, convex or concave nuclear shell surface with 167, 263 and 407 particles, as well as a resolution of 31 Å, 29 Å and 31 Å, respectively. In addition, the clean 886 particles were re-boxed at a pixel size of 6.78 Å to focus on a single chmA tetramer. This improved the resolution to 22 Å.

### RNA isolation, library preparation and sequencing

For RNA-seq, *E. coli* MC1000 cells were grown for 2-3 hr on 1% agarose and 25% LB plates in humidifier chambers and infected with Goslar. Samples were collected at 30 mpi from NT control and CRISPRi-ART knockdown cells, snap-frozen in liquid nitrogen, and stored at −20°C. RNA was isolated using an NEB Monarch Total RNA Miniprep Kit (T2010). The integrity and quality of RNA was analyzed using Agilent TapeStation 4150. For library preparation, 1 µg of total RNA was added in a nuclease-free PCR tube. Ribosomal RNA was removed and remaining RNA was purified using magnetic beads (NEBNext rRNA Depletion Kit (Bacteria) with RNA Sample Purification Beads, E7860L). cDNA was synthesized and libraries were prepared (NEBNext Ultra II RNA Library Prep Kit for Illumina, E7770S; NEBNext Multiplex Oligos for Illumina (Dual Index Primers Set 1), E7600S). The samples were sequenced using a NovaSeq X Plus Sequencing System (Illumina) platform and >20 million reads were collected per sample. Three independent biological replicates of each sample were sequenced.

### RNA sequencing analysis

RNA-seq reads were processed using an end-to-end analysis pipeline designed to enable direct comparison across datasets generated with different read lengths and sequencing depths. Raw paired-end reads were quality filtered and adapter trimmed using fastp [50](v1.3.6). Quality-control metrics, including read retention, mean read length, Q20, Q30, and GC content, were extracted from the fastp output for each sample and compiled into a master quality-control table (Table S6). Filtered reads were aligned to a combined reference genome containing both *E. coli* and bacteriophage Goslar sequences using Bowtie2/2.5.0 [51,52] with the -- very-sensitive-local alignment mode (Goslar NCBI ID NC_048170.1, *E. coli* NCBI ID NC_000913.3). Primary alignments were retained, sorted, indexed, and filtered using SAMtools [53,54] (v1.22.1). For downstream coverage analysis, alignments with mapping quality scores of MAPQ ≥10 were retained, and reads mapping specifically to the Goslar genome were extracted from the filtered BAM files. Primary mapping rates, properly paired reads, MAPQ ≥10 mapped reads, Goslar-mapped reads, and the fraction of MAPQ ≥10 reads mapping to Goslar are reported for each sample in Table S6. To account for differences in sequencing depth and read length among datasets, Goslar genome coverage was normalized using the total number of mapped Goslar bases rather than read counts alone. For each sample, the total number of Goslar-mapped bases was calculated from CI-GAR-aligned bases using SAMtools stats, and a scaling factor of 10^6^/(total Goslar-mapped bases) was applied during coverage-track generation. Coverage was calculated in 50-bp bins using bamCoverage from the deepTools suite(v3.5.6) [55]. Normalized coverage tracks were exported in BigWig (.bw) format using bedtools (v2.31.1) [56] and visualized using the Integrative Genomics Viewer (IGV_v2.19.8) [57], using identical display and Y-axis scaling within each comparison, with a fixed track height of 15; signal above this ceiling is clipped. Goslar transcriptional coverage was further quantified from MAPQ ≥10 Goslar-specific BAM files using SAMtools. Mean genome-wide RNA-seq depth was calculated as the total number of Goslar-mapped CIGAR bases divided by the Goslar genome length. Breadth of transcriptional coverage was calculated using samtools depth -aa and expressed as the percentage of Goslar genomic positions covered at ≥1× and ≥10× depth. Mean Goslar transcriptional depth and ≥1× and ≥10× coverage breadth for each library are reported in Table S6. Because the proportion of phage-derived RNA is expected to vary biologically across infection conditions, Goslar mapping fraction and transcriptional coverage were treated as descriptive measures of phage transcriptome representation rather than independent sequencing-quality pass/fail criteria. Unique reads mapped to the Goslar genome were quantified at the gene level using R package (v4.3.1) the featureCounts function in Rsubread (v2.14.2) to generate a count matrix [58] and Counts were assigned to Goslar CDS features using gene_id, with unstranded counting and paired-end fragment counting. The resulting raw count matrix was used directly as input to DESeq2 v1.40.2 [59]. DESeq2 models gene-level count data using a negative-binomial generalized linear model and estimates sample-specific size factors to account for differences in effective library depth. Wald tests were used to assess differential expression between conditions, and *P* values were adjusted for multiple testing using the Benjamini-Hochberg false-discovery-rate procedure. Cook’s-distance outlier handling was left at the DESeq2 default settings for all contrasts. Genes were considered differentially expressed when adjusted *P* < 0.05 and |log2 fold change| ≥ 2 (Table S5, Figure 4B, S11B). We further compared the transcriptional profiles of ChmA-depleted [7] and Rhc-depleted infections relative to their respective controls (Table S7).

For assessment of biological replicate similarity, DESeq2-normalized counts were transformed using the regularized logarithm (rlog) transformation. Pairwise Pearson correlation coefficients were calculated across samples using R’s cor function with method = “pearson”. A correlation-derived distance metric (1 − *r*) was used for hierarchical clustering of samples, and the resulting correlation matrix was visualized using the pheatmap package v1.0.12 [60] (Figure S11). Volcano plots were generated in R using ggplot2 v3.5.2 [61]. Horizontal and vertical dashed lines indicate the adjusted-*P* threshold of 0.05 and log2 fold-change thresholds of ±2 or ±1 depending upon the knockdowns. Genes were classified as significantly upregulated, significantly downregulated, or not significant according to these thresholds. Overlap between differentially expressed gene sets from different contrasts was calculated in R and visualized using the VennDiagram package v1.7.3 [62].

Sequencing depth differed between RNA-seq batches. ChmA-KD and control libraries published previously [7] were generated on the NovaSeq 6000 platform and showed lower mean Goslar coverage (~180–1,450×), whereas the Rhc knockdown and complementation libraries were generated on the NovaSeq X Plus platform and showed higher mean Goslar coverage (~1,850–8,180×). Consistent with this difference, replicate-to-replicate variability was modestly higher in the lower-depth batch (median CPM CV ~0.14) than in the higher-depth batch (median CV ~0.08). Differential-expression analysis was performed using DESeq2, which models replicate-level dispersion and normalizes for differences in library size. Sequencing-depth and coverage metrics for all samples are reported in Table S6.

## DATA AVAILABILITY

RNA-sequencing data, fluorescence microscopy data with larger example fields, cryo-ET data and 3-D reconstructions will be deposited to BioProject, Figshare, EMPIAR and EMDB respectively.

## ACKNOWLEDGEMENTS

We thank members of the Pogliano, Villa, and Corbett labs for helpful discussions. This work was supported by the HHMI Emerging Pathogens Initiative (to E.V., J.P., K.D.C., and K.P.), NIH grants R01 GM129245 (to J.P.) and R35 GM144121 (to K.D.C.), and NSF MRI grant DBI 1920374 (to E.V.). E.V. is a Howard Hughes Medical Institute Investigator; N.K. was supported by a Schmidt AI in Science Postdoctoral Fellowship. We thank the Translational Science Laboratory at the Florida State University College of Medicine for next-generation sequencing, and particularly Cynthia Vied for accommodating our urgent sequencing requests; the Biomolecular/Proteomics Mass Spectrometry Facility (BPMSF) at UC San Diego for mass spectrometry and Majid Ghassemian for running and analyzing the samples; the Institute for Genomic Medicine (IGM) Genomics Center at UC San Diego for reagents supporting library preparation; the UC San Diego cryo-EM facility, part of the Goeddel Family Technology Sandbox, and facility staff Mariusz Matyszewski and Inga Kuschnerus for their assistance; and UCSD Physics Computing director Kevin Smith and Brendan Dennis for computing support.

## AUTHOR CONTRIBUTIONS

Conceptualization, P.R., K.D.C., and E.V.; methodology, P.R., N.K., K.I., R.L., D.B., K.D.C., and E.V.; formal analysis, P.R., N.K., T.F., K.I., K.D.C., and E.V.; investigation, P.R., N.K., R.L., M.L., T.F., S.M.B., C.R., L.Y.H., and M.N.; resources, E.G.A. and C.J.M.; data curation, P.R., N.K., T.F., and K.I.; visualization, P.R., N.K., K.D.C., M.R., and T.F.; supervision, K.D.C., J.P., and E.V.; writing of original draft, P.R., N.K., K.D.C., and E.V.; writing, review & editing, all authors.

## SUPPLEMENTAL FIGURES

**Figure S1.**
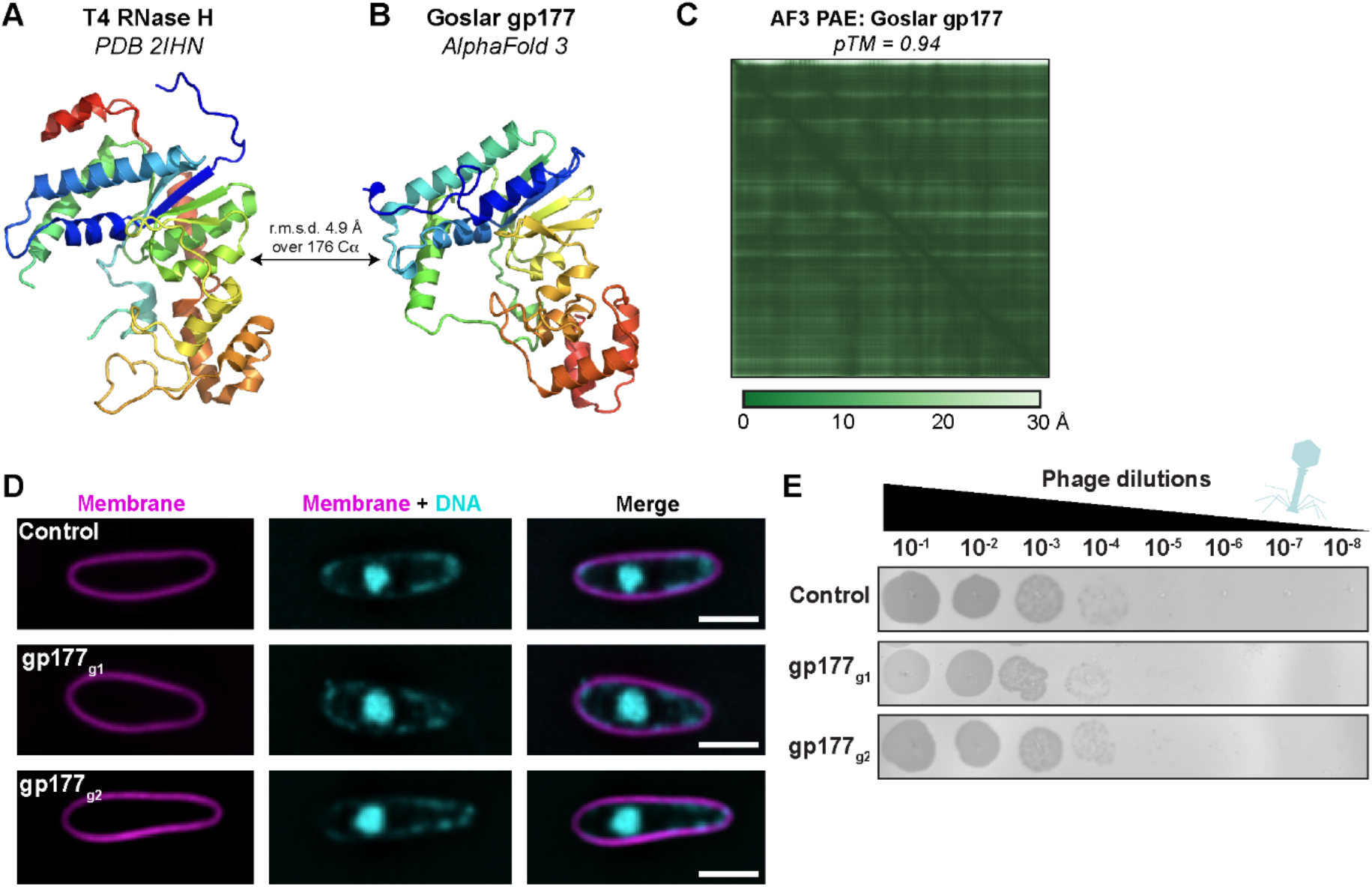
gp177 is dispensable for Goslar phage replication. **(A)** Crystal structure of T4 phage RNase H (PDB 2IHN) [63], colored as a rainbow from N-terminus (blue) to C-terminus (red). **(B)** AlphaFold 3 predicted structure of Goslar gp177, colored as a rainbow from N-terminus (blue) to C-terminus (red). **(C)** AlphaFold 3 predicted aligned error (PAE) plot for the structure prediction shown in panel (B). **(D)** Fluorescence microscopy analysis of non-targetting control (NT control) and gp177 knockdown infections. Cells were stained with FM4-64 to label membranes (magenta) and DAPI to visualize DNA (cyan). Scale bars, 2 µm. **(E)** Spot titer assays comparing phage progeny production in non-targeting control (NT) and gp177 knockdown strains using two independent guide RNAs (g1 and g2). The indicated serial dilutions of Goslar were spotted onto *E. coli* lawns.

**Figure S2.**
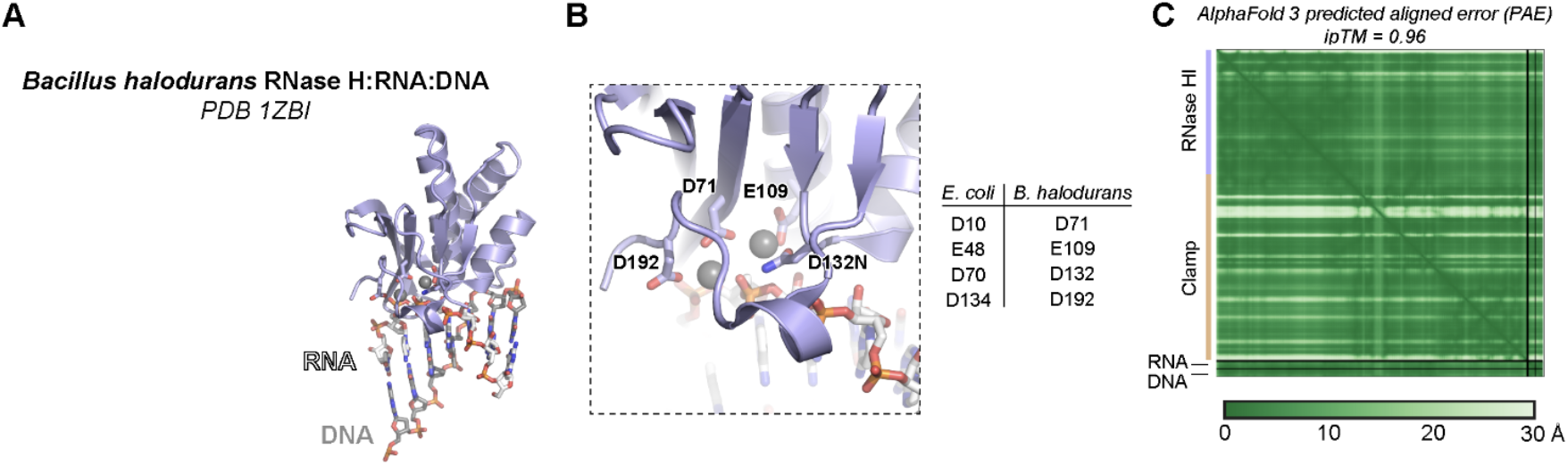
Rhc has a predicted RNase HI domain. **(A)** Structure of *B. halodurans* RNase H bound to an RNA-DNA hybrid (PDB ID 1ZBI) [64]. **(B)** Closeup of the *B. halodurans* RNase H catalytic site, with conserved catalytic residues shown as sticks and two Mg^2+^ ions shown as spheres. *Right:* Equivalent active site residues of *E. coli* RNase H and *B. halodurans* RNase H. **(C)** AlphaFold 3 predicted aligned error (PAE) plot for the prediction shown in panel (

**Figure S3.**
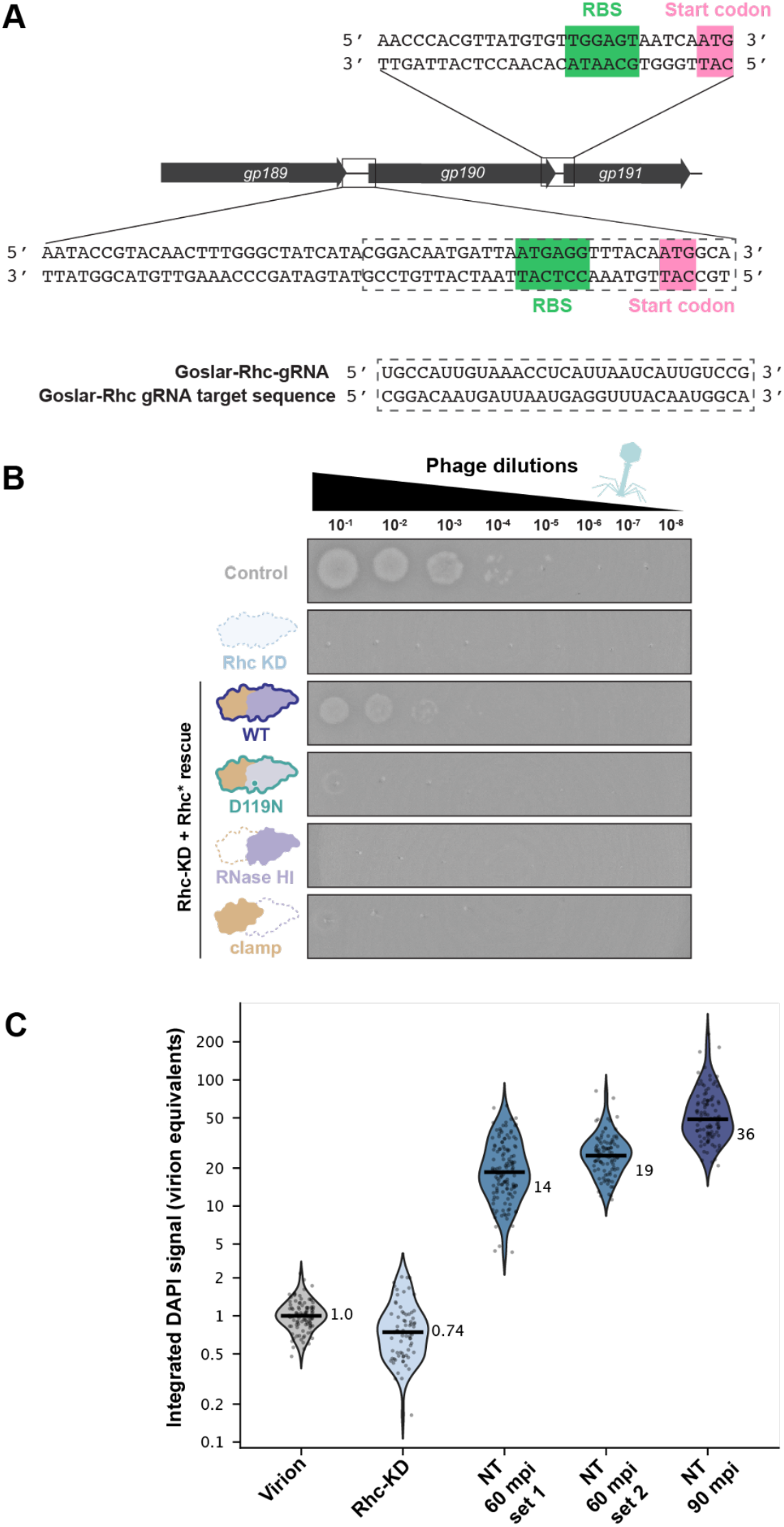
Goslar Rhc CRISPRi-ART knockdown strategy and phenotype. **(A)** Schematic showing the genomic context of Goslar *gp190* (*Rhc*) and CRISPRi-ART guide RNA. Expanded views highlight the *gp190* 5^′^ untranslated and coding regions, indicating the likely ribosome-binding site (RBS; green) and start codon (magenta). The CRISPRi-ART guide RNA (gp190_gRNA) was designed to target the translation initiation region. **(B)** Representative spot titer assay of infectious phage production under gp190/Rhc knockdown and complementation by wild type (WT) or catalytic dead (D119N) CRISPRi-ART-resistant Rhc (Rhc*), or the isolated RNase H or clamp domains. Goslar lysates were pre-diluted 1:400, followed by 10-fold serial dilution and spotting on bacterial lawns. Dilution labels indicate the serial dilution factor applied after the initial 1:400 pre-dilution. **(C)** Preliminary analysis of DAPI signal per nucleus, expressed in virion equivalents (integrated DAPI signal relative to the median of free virions; n = 99 virions, 72 Rhc-KD nuclei at 60 mpi, and non-targeting control (NT) nuclei: 133 (set 1) and 100 (set 2) at 60 mpi and 101 at 90 mpi). Violins show kernel density estimates of log10(value), bars show medians, and dots show individual objects.

**Figure S4.**
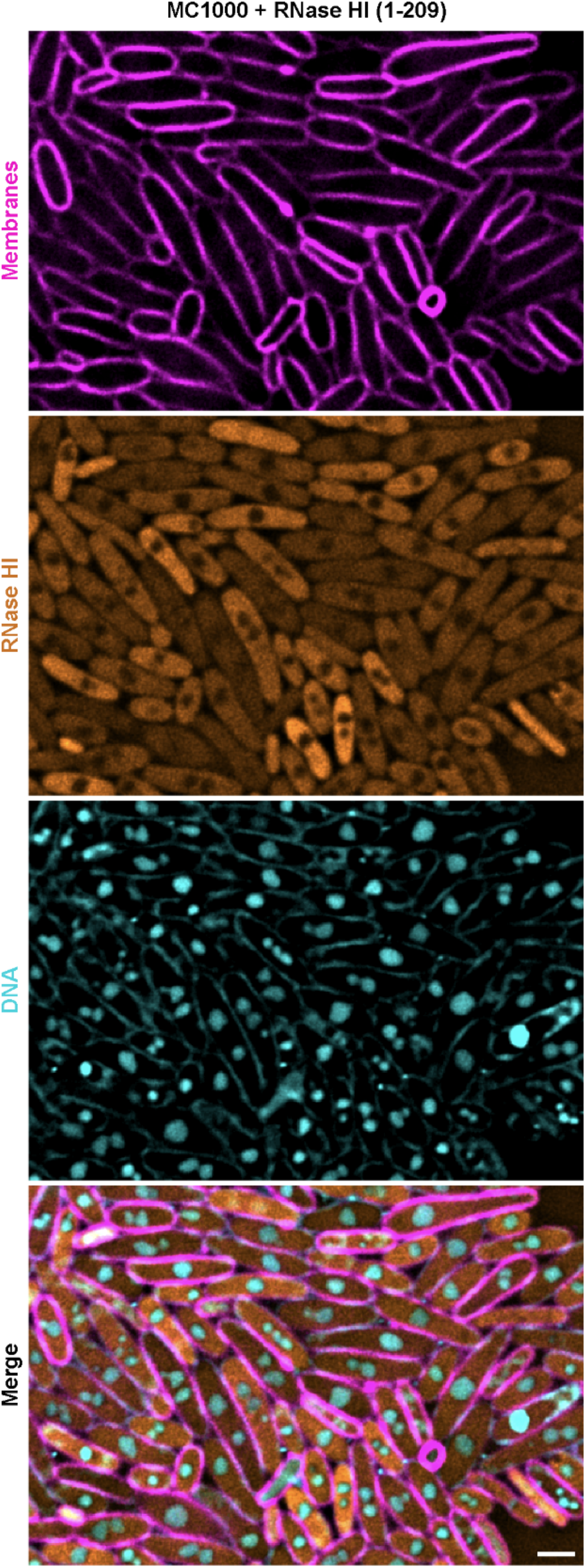
Localization of GFP-Rhc* RNase HI domain Goslar-infected *E. coli* MC1000. Representative fluorescence images of *E. coli* MC1000 expressing GFP-Rhc* RNase HI domain (residues 1-209). Cell membranes were labeled with FM4-64 (magenta), GFP-Rhc* is shown in orange, and DNA was stained with DAPI (cyan). Scale bar = 2 μm.

**Figure S5.**
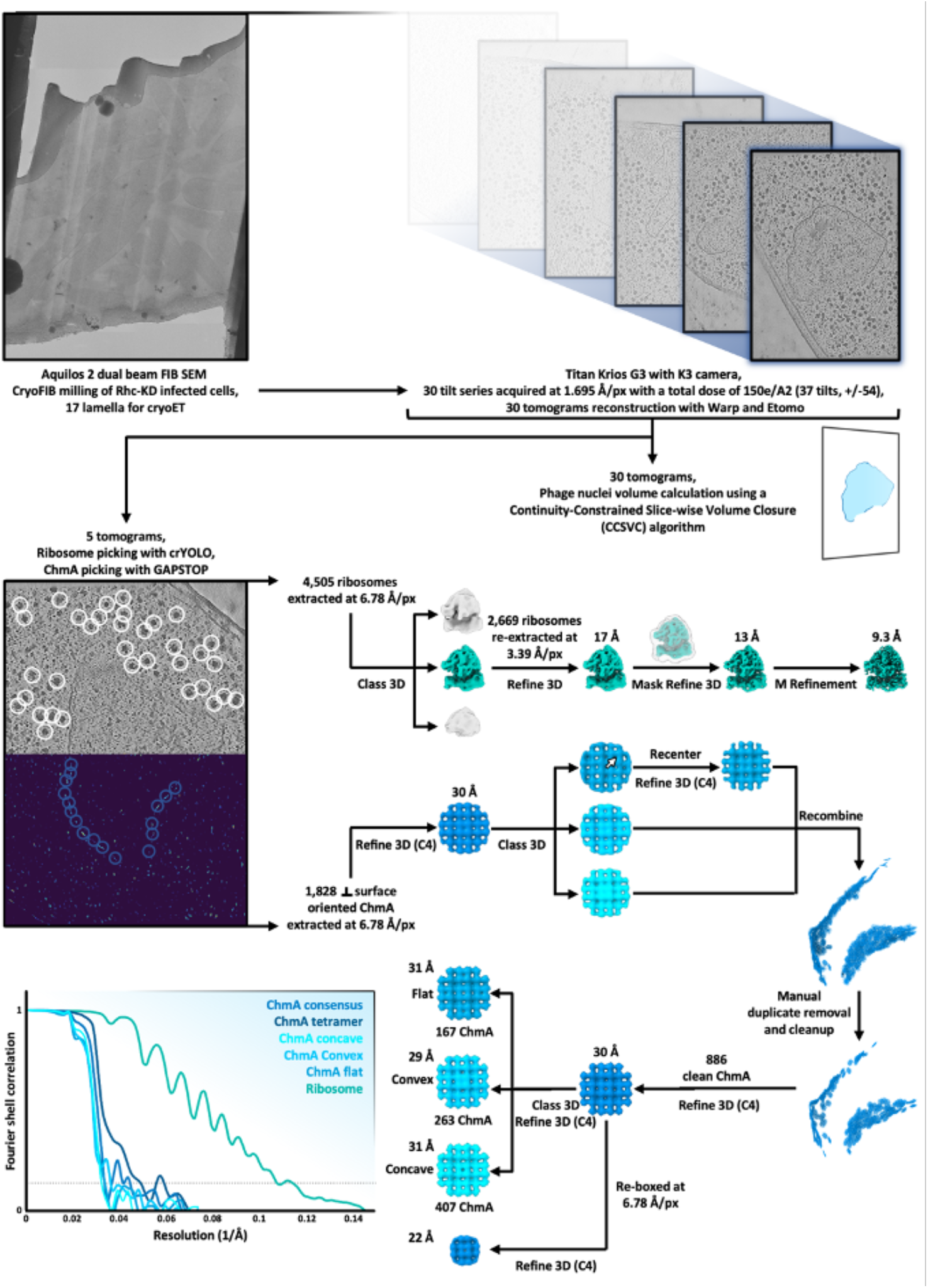
*In situ* cryoET data acquisition and subtomogram analysis of bacterial ribosomes and phage chmA in Goslar-infected Rhc-KD cells. Rhc-KD cells were cryoFIB-milled 60 min after Goslar infection, yielding 17 lamellae from which 30 tilt series (±54°, 150 e−/Å^2^) were acquired on a Titan Krios G3 equipped with a K3 detector. Tomograms were reconstructed using Warp and Etomo, and phage nuclei volumes were quantified using the CCSVC algorithm. Ribosomes and chmA were picked from five tomograms using crYOLO and GAPSTOP, respectively. Ribosome sub-tomogram analysis yielded a final resolution of 9.3 Å following refinement in the Relion/Warp/M pipeline. ChmA tetramers were refined with C4 symmetry and 3D classified to identify picking outliers. Duplicates and orientation outliers were then manually removed or corrected. Subsequent additional 3D classification of the clean dataset and focused 3D refinement resolved flat, convex, and concave shell conformations at 31-22 Å resolution, based on the FSC 0.143 criterion (grey dotted line).

**Figure S6.**
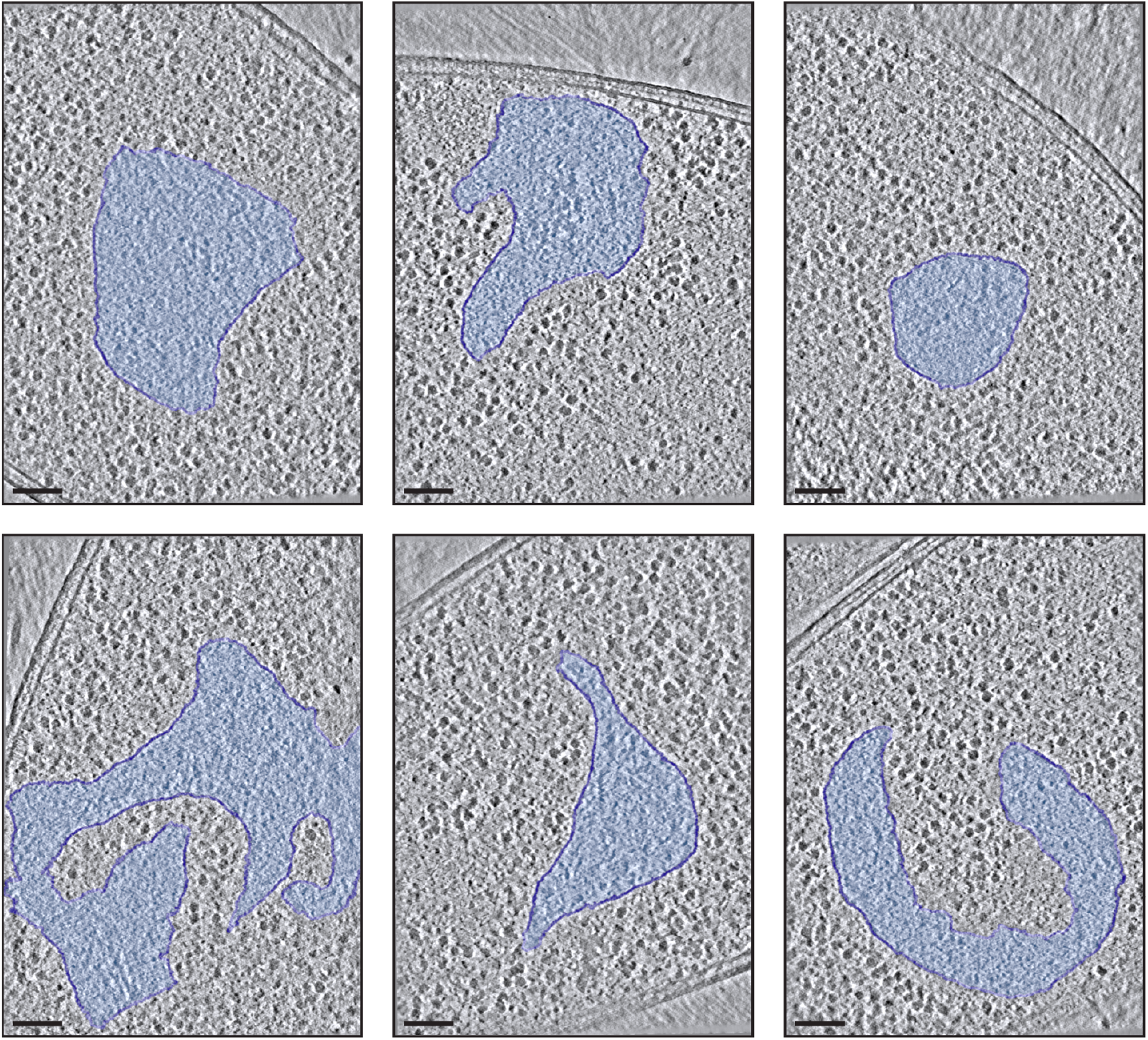
*In situ* cryoET of Rhc KD phage nuclei. Tomographic slices of cryoFIB milled *E. coli* cells show the different shapes of the nucleus 60 min after phage infection. Scale bar = 0.1 µm.

**Figure S7.**
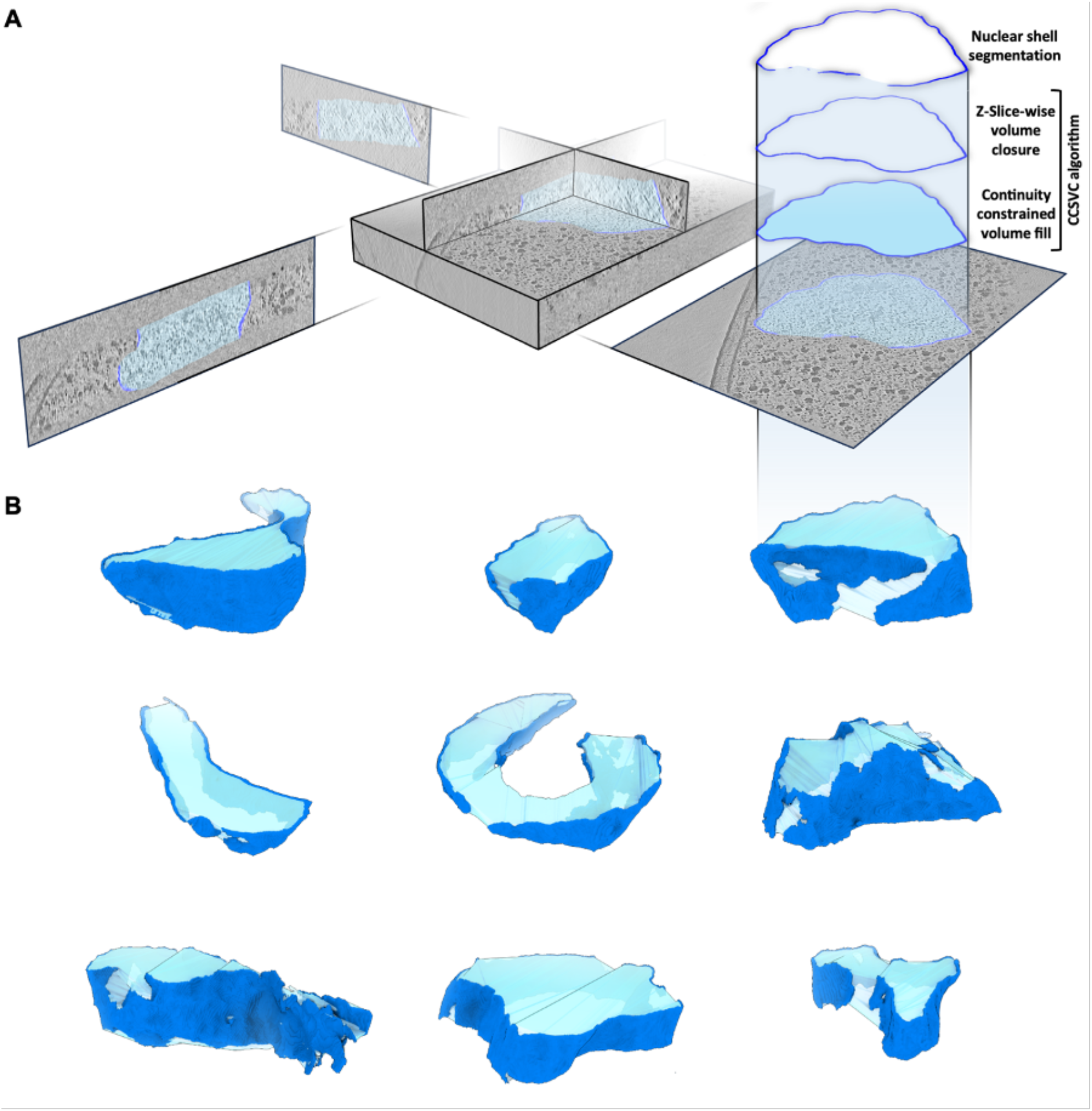
Phage nuclei volume calculation using the Continuity-Constrained, Slice-wise Volume Closure (CCSVC) algorithm. **(A)** Reconstructed tomograms of Rhc-KD cells were segmented to obtain a cryoEM density of the embedded phage nuclei. The segmentation was then converted into a binary mask and skeletonized to define endpoints for each z-slice. Endpoint pairs were connected by straight voxel paths to close boundaries in the preliminary fragmented segmentation of the phage nucleus. A closed binary mask allowed for subsequent filling of the cross-sectional area in each z-slice. **(B)** The CCSVC algorithm allowed for the generation of a three-dimensional binary volume enclosed by the phage nuclei. Generated volumes (transparent cyan) with their surrounding fragmented nuclear shell segmentation (blue) are shown for a subset of tomograms.

**Figure S8.**
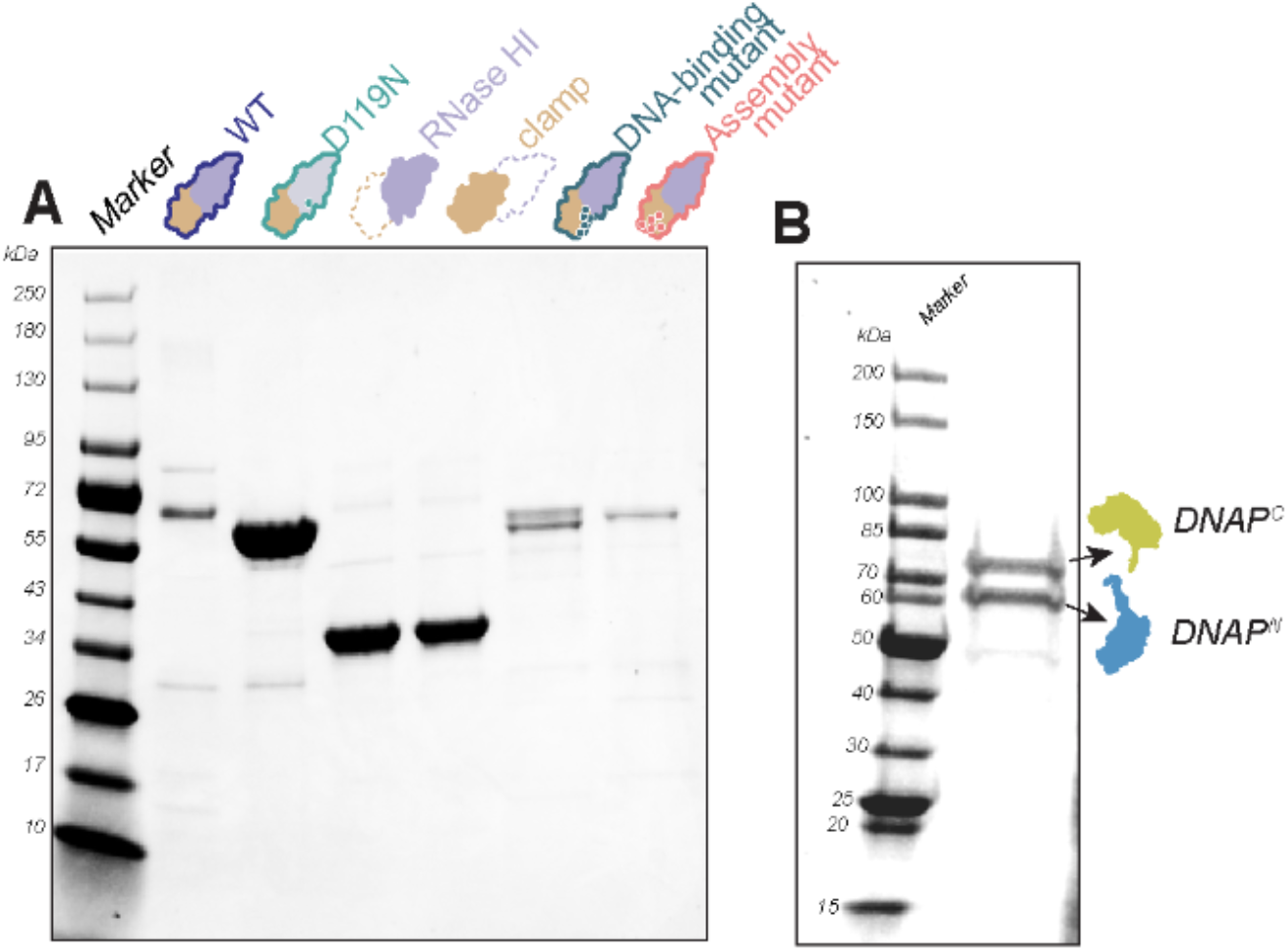
Purification of Rhc variants and the Goslar DNA polymerase complex. **(A)** Coomassie-stained SDS-PAGE showing purified WT, the catalytic-dead D119N mutant, the isolated RNase HI domain, the isolated clamp domain, the DNA-binding mutant, and the assembly mutant. Molecular-weight markers are indicated in kDa. **(B)** Coomassie-stained SDS-PAGE of the purified Goslar DNAP complex containing DNAP-C and DNAP-N. The positions of DNAP^C^ and DNAP^N^ are indicated by arrows. Molecular-weight markers are shown in kDa.

**Figure S9.**
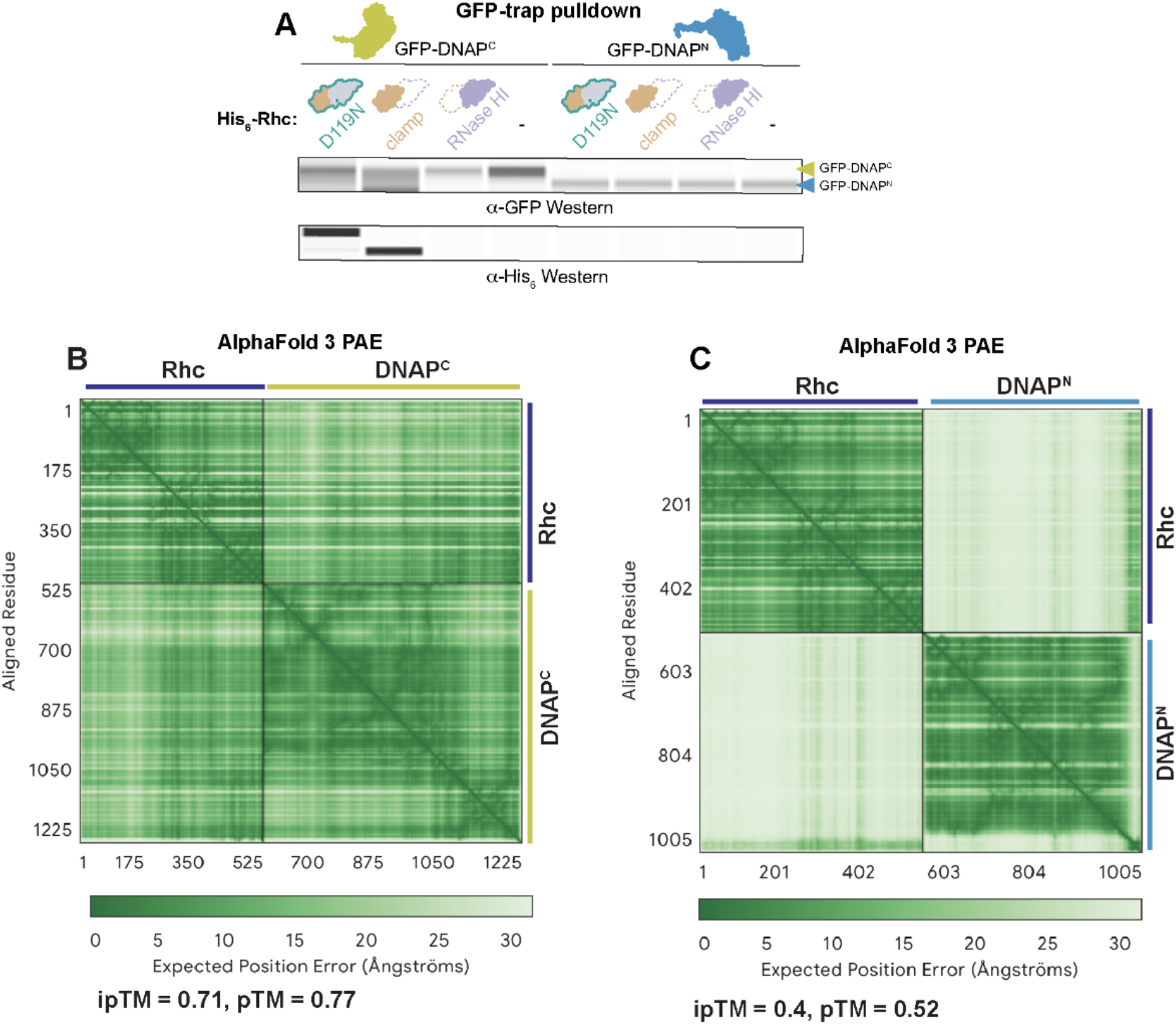
Rhc is a DNA polymerase subunit. **(A)** GFP affinity pull-down assay examining interactions between phage DNA polymerase subunits (DNAP^C^ and DNAP^N^) and Rhc. Elution fractions were visualized by Jess capillary western blotting against GFP (top) and His-6 (bottom). **(B)** PAE plot for the Rhc-DNAP^C^ complex. **(C)** PAE plot for the Rhc-DNAP^N^ complex.

**Figure S10.**
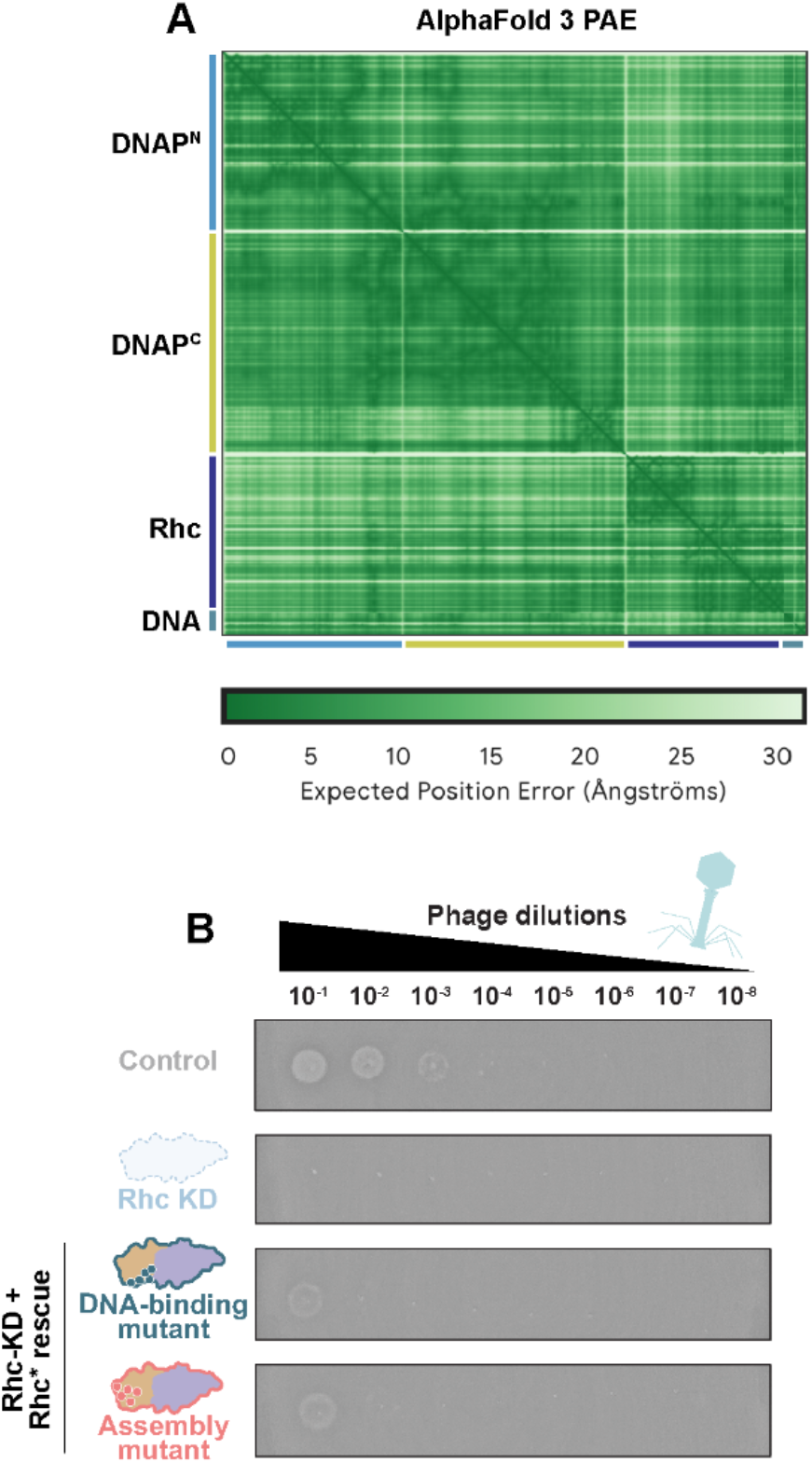
Rhc association with the DNA polymerase complex and requirement for productive infection. **(A)** AlphaFold3 predicted aligned error (PAE) plot for the complex comprising DNAP^N^, DNAP^C^, Rhc, and DNA. Lower PAE values indicate higher confidence in the relative positioning of the corresponding regions within the predicted complex. **(B)** Spot-titer assay showing productive Goslar infection under the indicated conditions. Serial 10-fold phage dilutions (10^−1^-10^−8^) were spotted on control cells, Rhc-knockdown (KD) cells, or Rhc-KD cells complemented with the indicated Rhc* DNA-binding or assembly mutants. Rhc depletion strongly impaired plaque formation, and neither mutant restored productive infection.

**Figure S11.**
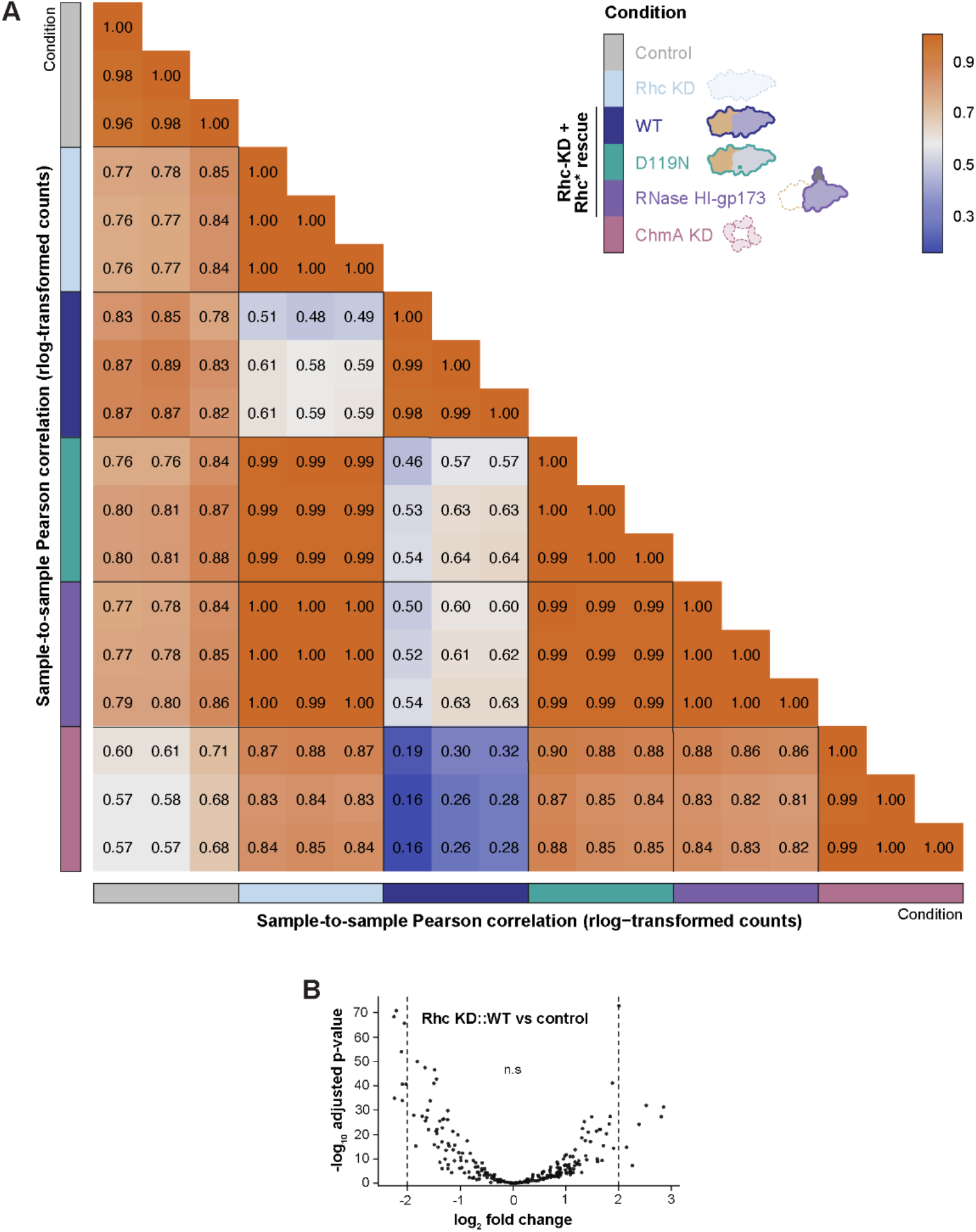
Sample-sample Pearson correlation analysis and Rhc complementation with WT Rhc. **(A)** Heatmap showing pairwise Pearson correlation coefficients calculated from rlog-transformed RNA-seq expression values across all samples, including non-targeting controls (Control), Rhc knockdown (Rhc KD), ChmA knockdown (ChmA KD), and RNaseH knockdown complemented with WT, RNase H catalytic mutant (D119N), or RNase H domain fused to gp173 (gp173 RNase H). Samples are hierarchically clustered based on correlation-derived distances, and correlation strength is indicated by the color scale (purple to yellow, low to high). Biological replicates cluster tightly within each condition, demonstrating high reproducibility. Full-length Rhc complementation clusters closely with non-targeting controls, whereas Rhc KD, catalytic mutant (D119N), and NTD-Taileboi conditions cluster separately, indicating that catalytic activity and full-length Rhc architecture are required to restore the global transcriptional state. Colored annotation bars indicate experimental conditions. **(B)** Volcano plot of RNA-seq differential expression analysis comparing Rhc knock-down (KD) complemented with wild-type (WT) Rhc with control. Each point represents a phage gene. The x-axis shows the log_2_ fold change, and the y-axis shows the −log_10_ adjusted *P* value. Dashed vertical lines indicate the fold-change threshold (±2), and the horizontal dashed line indicates the adjusted *P* value significance threshold (0.05). The majority of genes are not significantly differentially expressed, demonstrating that Rhc depletion does not broadly alter phage gene expression.

**Figure S12.**
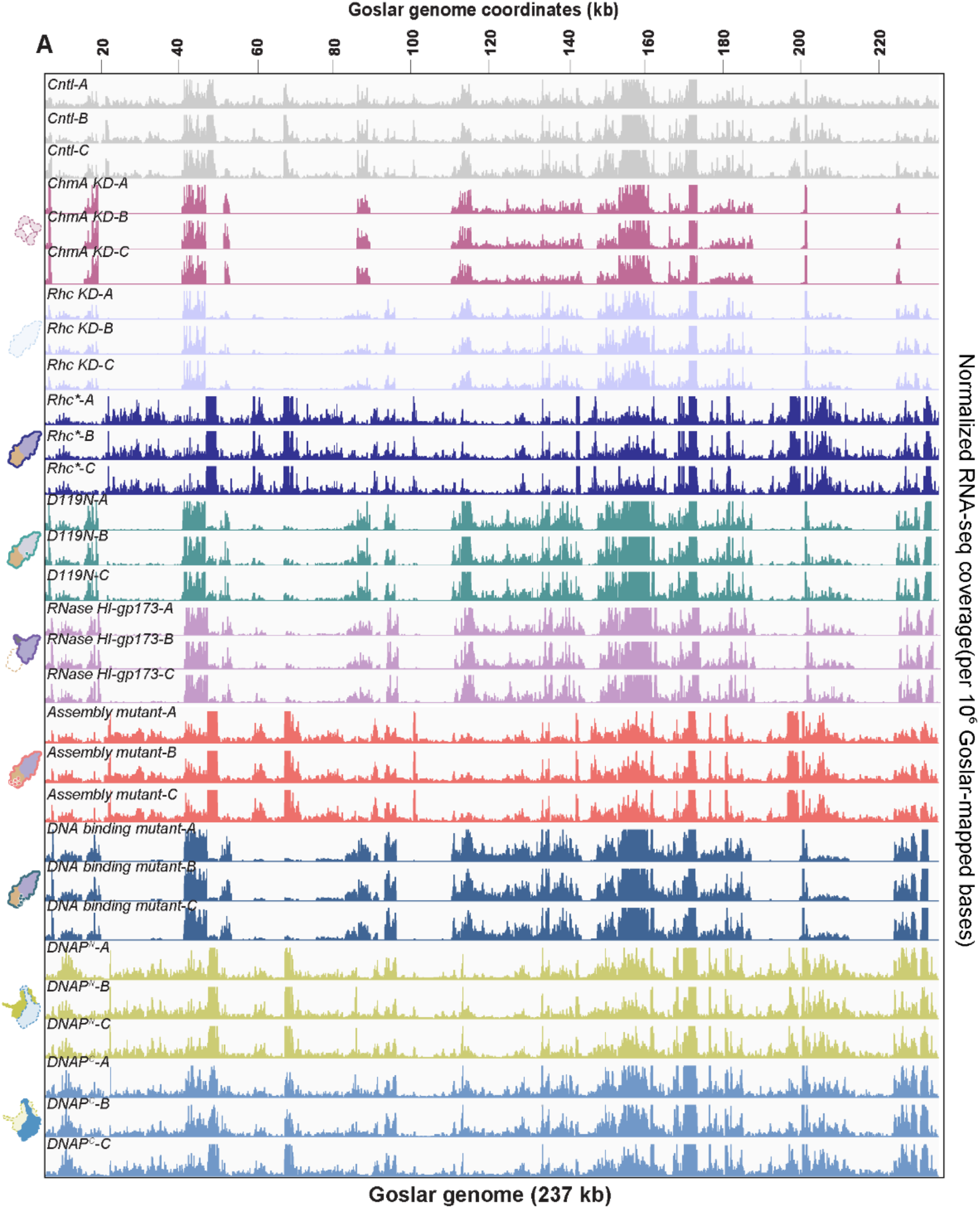
Transcriptomics profile of Rhc Knockdown and various complementations. Genome-wide RNA-seq coverage across the 237-kb Goslar genome under the indicated infection conditions. RNA-seq reads were aligned to a combined *E. coli*-Goslar reference, filtered to retain primary alignments with MAPQ ≥10 (mapping quality score), and reads mapping to the Goslar genome were extracted for visualization. For each sample, Goslar coverage was normalized by the total number of mapped Goslar bases using a scale factor of **1**0^6^/(total Goslar-mapped bases), thereby accounting for differences in sequencing depth and read length between datasets. Normalized coverage tracks were exported as BigWig (.bw) files and visualized in IGV. Three biological replicates are shown for each condition, including the non-targeting control (Cntl), ChmA knockdown (ChmA KD), Rhc knockdown (Rhc KD), the catalytically inactive Rhc D119N mutant, RNase H I tagged to gp173 perturbation, DNA binding mutant,and depletion of the N- and C-terminal subunits of DNA polymerase subunits (DNAP^N^ and DNAP^C^, respectively). IGV tracks were displayed at a fixed track height of 15. Goslar genome coordinates are shown above the tracks.

**Figure S13.**
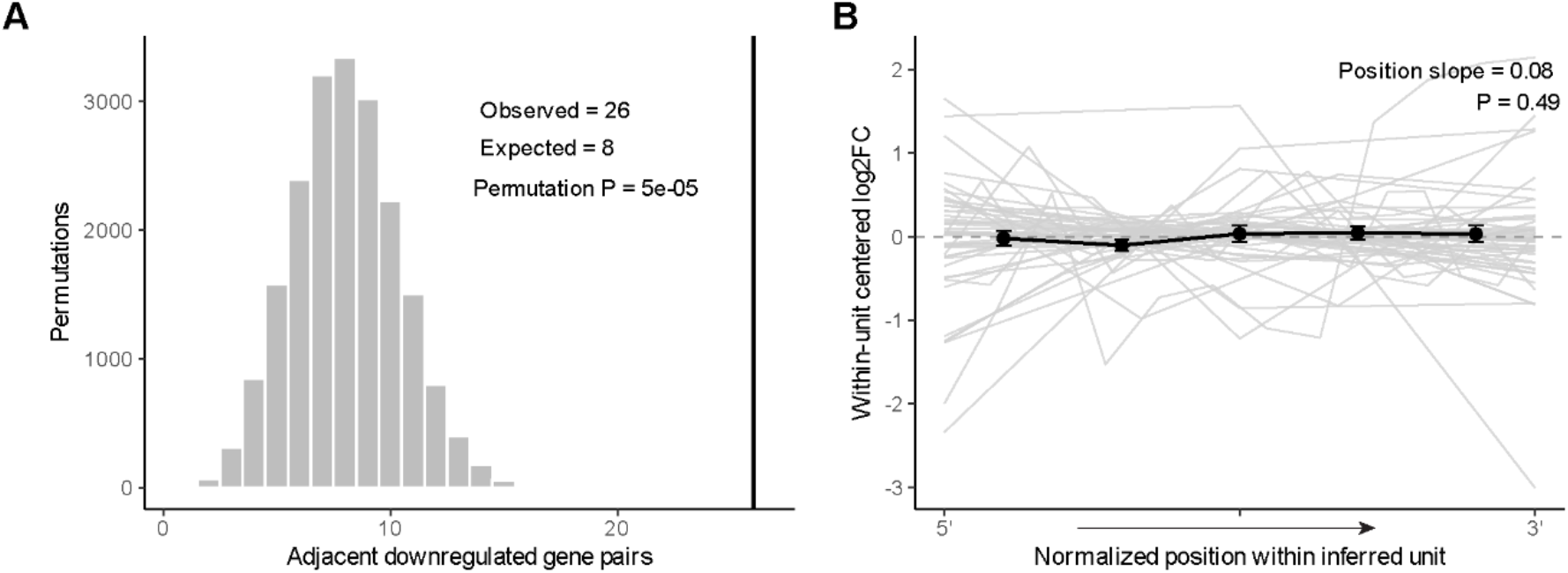
Rhc-dependent transcriptional defects are non-randomly clustered but do not show progressive 5′ to 3′ loss within putative transcriptional units. **(A)** Permutation analysis of genomic clustering of Rhc-dependent genes. Rhc-dependent downregulated genes were defined as genes with adjusted *P* < 0.05 and log_2_ fold change ≤ −2 in Rhc knockdown relative to control. The observed number of adjacent downregulated gene pairs was compared with the distribution obtained from 20,000 random permutations while preserving the number of downregulated genes on each strand. The observed dataset contained 26 adjacent downregulated gene pairs, compared with a random expectation of approximately 8 pairs. The vertical line indicates the observed value. The observed clustering exceeded that obtained in the randomized datasets (empirical permutation *P* = 5 × 10^−5^), demonstrating that Rhc-dependent downregulated genes are significantly more spatially clustered than expected by chance. **(B)** Analysis of transcriptional position within inferred transcriptional units. Inferred units were defined as contiguous genes on the same strand separated by ≤100 bp. Genes within each unit were ordered according to their true 5′ to 3′ transcriptional direction, with minus-strand units reversed relative to genomic coordinate order. Gene position within each unit was normalized from 0 at the 5′ end to 1 at the 3′ end, allowing transcriptional units of different sizes to be compared on a common scale. To isolate within-unit positional effects from differences in overall expression between units, the mean log_2_ fold change of each unit was subtracted from the log_2_ fold change of each constituent gene, generating a within-unit centered log_2_ fold change. Gray lines represent individual inferred transcriptional units, and black points and lines summarize the mean positional trend. Linear regression of centered log_2_ fold change against normalized 5′-to-3′ position yielded a slope of 0.08 (*P* = 0.49), indicating no significant progressive increase in downregulation toward the 3′ ends of inferred transcriptional units. Thus, although Rhc-dependent genes are strongly clustered across the genome, the transcriptional defect is not consistent with a simple genome-wide model of progressively reduced RNA polymerase processivity from 5′ to 3′.

**Figure S14.**
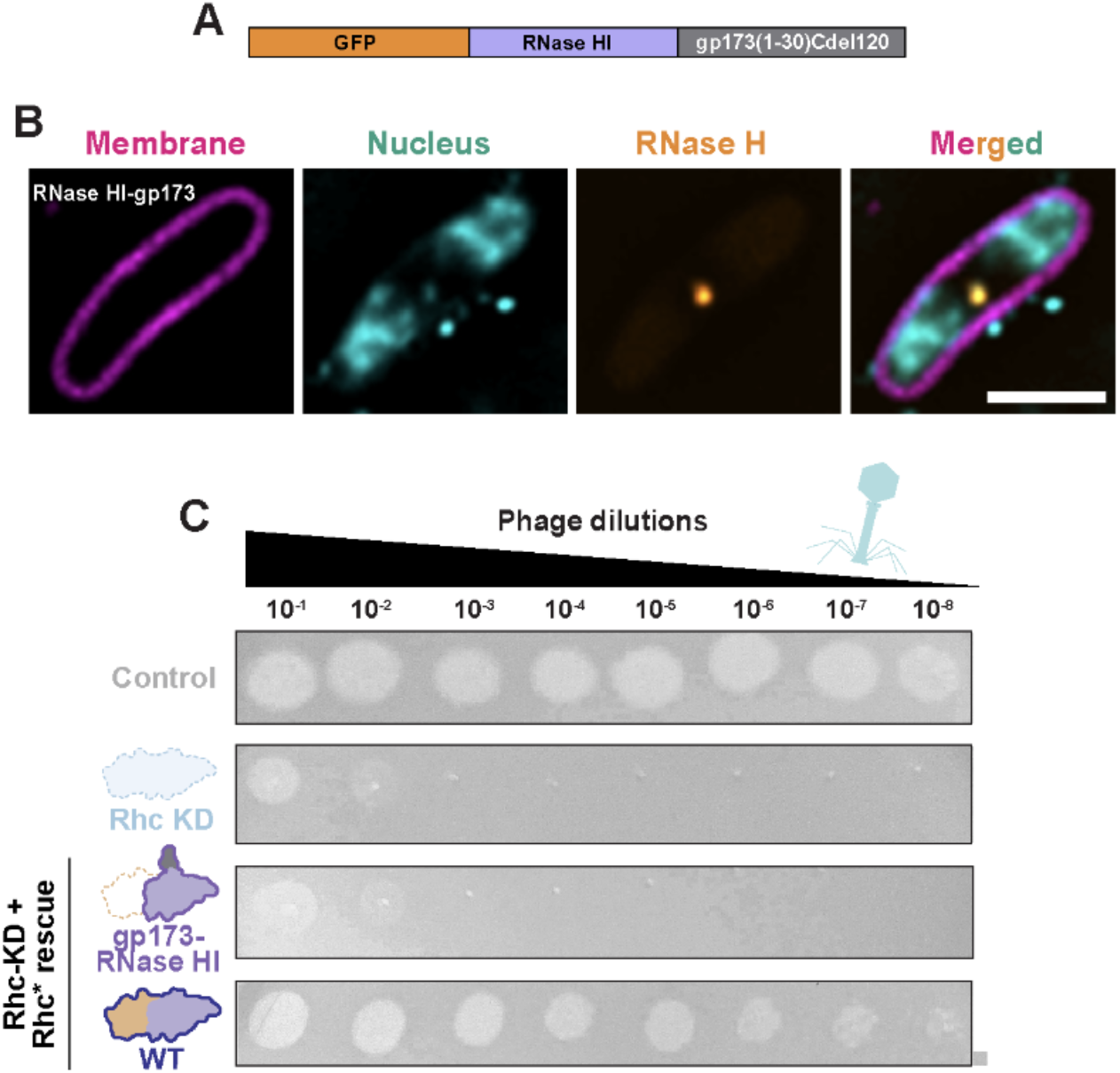
Forced nuclear targeting of the N-terminal RNase H domain is insufficient to rescue phage replication. **(A)** Schematic of the fusion construct used to force nuclear targeting of the RNaseH N-terminal domain (NTD). GFP-tagged RNaseH-NTD was fused to residues 1-30 of gp173 (Taileboi), with deletion of residues 1-120, a construct previously shown to localize to the phage nucleus. **(B)** Fluorescence microscopy showing forced localization of the N-terminal RNase H domain to the nascent nucleus. **(C)** Spot titer assay showing that forced nuclear localization of the N-terminal RNase H domain fails to rescue phage propagation and lysis.

**Figure S15.**
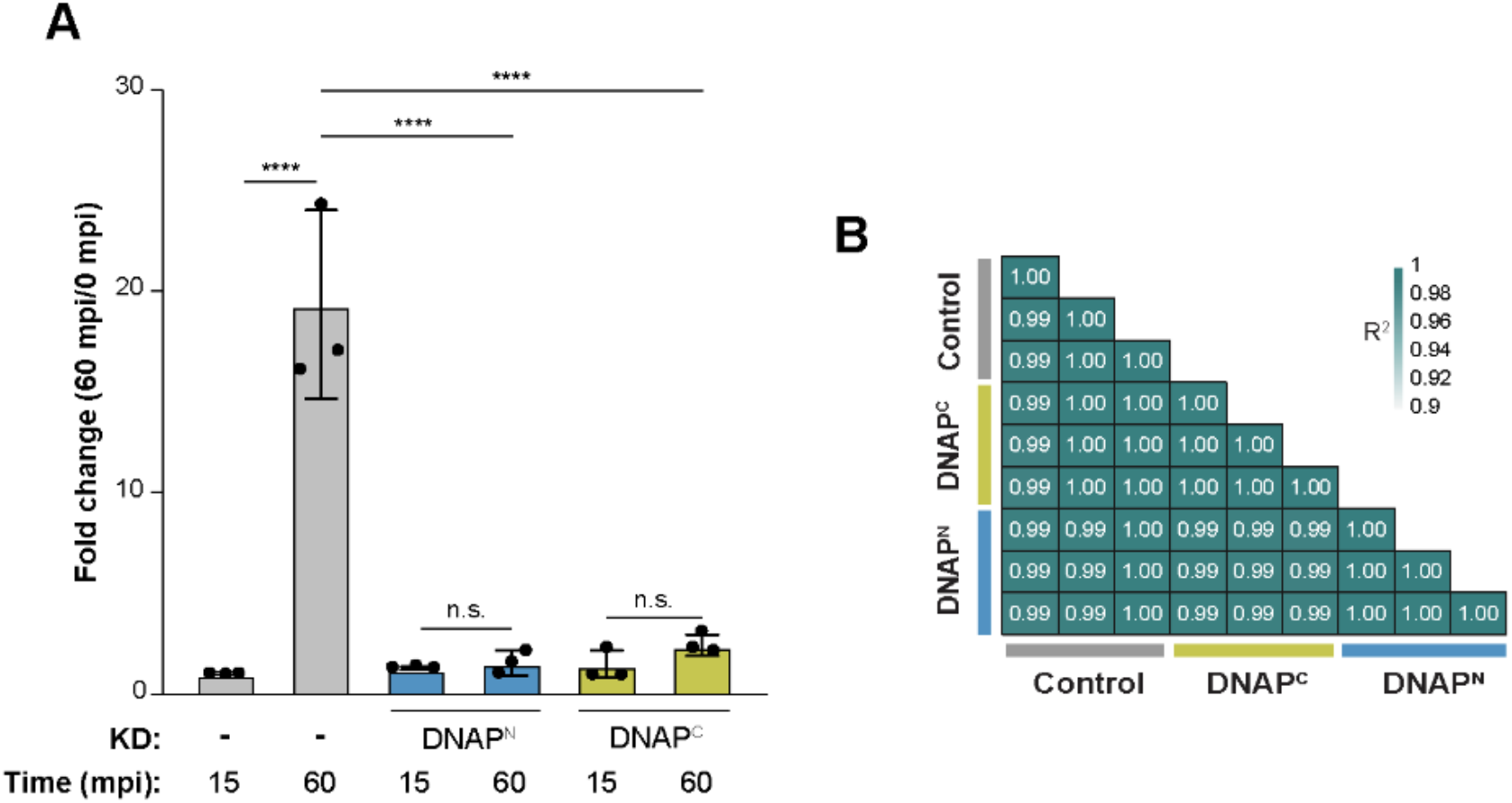
DNA polymerase knockdown disrupts phage genome replication while largely preserving the transcriptional program. **(A)**. Measured at indicated time points and normalized by 0 mpi. Control infections show robust genome amplification at 60 mpi, whereas DNAP knock-downs (DNAP^N^ and DNAP^C^ KD) infections fail to support DNA replication. Bars represent mean ± s.e.m. (n = 3 biological replicates). Statistical significance was assessed by one-way ANOVA followed by Šídák’s multiple comparisons test (α = 0.05). Comparisons of control versus KDs conditions were highly significant (****P < 0.0001). **(B)**. Pairwise correlation matrix comparing genome-wide phage RNA-seq profiles from the non-targeting control, DNAP^C^-depleted, and DNAP^N^-depleted infections. Individual tiles represent pairwise coefficients of determination, R-squared. The high correlations among conditions indicate that depletion of either DNA polymerase subunit has little effect on the overall phage transcriptional profile.

## SUPPLEMENTAL TABLES

**Table S1.** Goslar Rhc C-terminal domain DALI hits. Top 25 hits from the PDB25 subset. Light green highlight indicates sliding clamp proteins

| PDB-Chain | Z score | Calpha rmsd | Alignment length | Uniprot ID | Description |
| --- | --- | --- | --- | --- | --- |
| 3ggr-B | 7.1 | 4.7 | 179 | O60921 | H. sapiens HUS1 |
| 7st9-G | 5.9 | 4 | 166 | Q08949 | S. cerevisiae Ddc1 |
| 3a1j-A | 5.7 | 4 | 156 | Q99638 | H. sapiens RAD9A |
| 7st9-H | 5.3 | 4.8 | 180 | Q02574 | S. cerevisiae Mec3 |
| 7ype-A | 5.3 | 4.3 | 147 | A0A0A1E3R5 | African swine fever virus E301R |
| 2awa-B | 4.9 | 4.5 | 159 | O06672 | Streptococcus pneumoniae TIGR4 (DNA Pol III beta chain) |
| 3hi8-A | 4.7 | 4.5 | 148 | D0VWY8 | Haloferax volcanii PCNA |
| 6drt-A | 4.7 | 3.8 | 143 | P04525 | phage T4 gp45 |
| 4ftx-A | 4.5 | 3.8 | 87 | P38247 | S. cerevisiae Slm4 |
| 3ggr-C | 4.4 | 3.6 | 148 | O60671 | H. sapiens RAD1 |
| 1tu1-A | 4.3 | 4.9 | 84 | Q9I738 | P. aeruginosa PA94 |
| 6y2p-A | 4.3 | 5.4 | 71 | P52129 | E. coli RnlA toxin |
| 7wmp-a | 4.3 | 3.2 | 75 | I7HHN4 | Helicobacter phage KHP30 portal |
| 6e8a-A | 4.2 | 3.9 | 81 | Q7CPJ3 | Salmonella enterica DcrB |
| 5xqi-B | 4.2 | 4.8 | 51 | Q9GZN7 | H. sapiens Rogdi |
| 7kzn-I | 4.2 | 3.9 | 67 | Q9SWQ6 | C. reinhardtii dynein light chain roadblock LC7a |
| 6aig-A | 4.2 | 5.5 | 154 | Q9YFT8 | Aeropyrum pernix K1 PCNA |
| 2du7-A | 4 | 6 | 97 | Q59054 | M. jannaschii O-phosphoseryl-tRNA synthetase |
| 8yqg-B | 4 | 3.2 | 70 | A0A5B9DBT7 | Promethearchaeum syntrophicum longin domain protein |
| 8es4-E | 4 | 11.7 | 67 | A0A482JLu9 | Shigella phage Buco gp40 |
| 6n90-A | 4 | 3.4 | 70 | A9CID9 | Agrobacterium tumefaciens YoeB |
| 4kc5-A | 3.9 | 7.9 | 114 | A1KQS1 | Mycetohabitans rhizoxinica RhiE |
| 6e8e-A | 3.8 | 4.5 | 156 | P0A988 | E. coli beta sliding clamp DnaN |
| 6xrb-A | 3.8 | 7.5 | 92 | A0A0H3NHS6 | Salmonella enterica SciW |
| 5iwd-A | 3.8 | 4.3 | 134 | P16790 | Human cytomegalovirus UL44 |

**Table S2.** Goslar GFP-Rhc IP-MS results. Pink shading indicates an RNA polymerase subunit; green shading indicates a DNA polymerase subunit.

| NCBI # | Name | Sequence coverage (%) | # peptides | Found in GFP control? |
| --- | --- | --- | --- | --- |
| YP_009820929.1 | putative ribonuclease HI HOV27_gp190 | 81 | 77 | No |
| YP_009820859.1 | Nuclear import factor PicA HOV27_gp174 | 25 | 9 | Yes |
| YP_009820893.1 | hypothetical protein HOV27_gp226 | 30 | 5 | Yes |

| NCBI # | Name | Sequence cover-<br>age (%) | # peptides | Found in GFP<br>control? |
| --- | --- | --- | --- | --- |
| YP_009820873.1 | Nuclear shell protein ChmA HOV27_gp246 | 26 | 10 | Yes |
| YP_009820726.1 | Major capsid protein HOV27_gp041 | 11 | 6 | Yes |
| YP_009820905.1 | hypothetical protein HOV27_gp214 | 46 | 5 | Yes |
| YP_009820723.1 | hypothetical protein HOV27_gp038 | 8 | 5 | Yes |
| YP_009820856.1 | nvRNAP subunit HOV27_gp171 | 9 | 4 | Yes |
| YP_009820753.1 | DNA Polymerase HOV27_gp068 | 8 | 3 | No |
| YP_009820708.1 | hypothetical protein HOV27_gp023 | 25 | 2 | No |
| YP_009820724.1 | nvRNAP subunit HOV27_gp039 | 7 | 2 | No |
| YP_009820755.1 | hypothetical protein HOV27_gp070 | 22 | 2 | Yes |
| YP_009820782.1 | hypothetical protein HOV27_gp097 | 35 | 2 | No |
| YP_009820861.1 | Nuclear shell-associated protein ChmC HOV27_gp176 | 13 | 2 | No |
| YP_009820743.1 | hypothetical protein HOV27_gp058 | 7 | 2 | Yes |
| YP_009820689.1 | hypothetical protein HOV27_gp004 | 14 | 2 | Yes |
| YP_009820704.1 | hypothetical protein HOV27_gp019 | 16 | 1 | Yes |
| YP_009820788.1 | hypothetical protein HOV27_gp103 | 17 | 1 | Yes |
| YP_009820850.1 | nvRNAP subunit HOV27_gp165 | 2 | 1 | No |
| YP_009820869.1 | hypothetical protein HOV27_gp184 | 8 | 1 | No |
| YP_009820882.1 | hypothetical protein HOV27_gp237 | 4 | 1 | No |
| YP_009820794.1 | hypothetical protein HOV27_gp109 | 9 | 1 | No |
| YP_009820746.1 | hypothetical protein HOV27_gp061 | 4 | 1 | No |
| YP_009820789.1 | hypothetical protein HOV27_gp104 | 17 | 1 | Yes |
| YP_009820694.1 | Putative prohead protease HOV27_gp009 | 4 | 1 | No |
| YP_009820756.1 | hypothetical protein HOV27_gp071 | 9 | 1 | No |
| YP_009820931.1 | hypothetical protein HOV27_gp188 | 3 | 1 | No |
| YP_009820860.1 | nvRNAP subunit HOV27_gp175 | 2 | 1 | No |
| YP_009820811.1 | hypothetical protein HOV27_gp126 | 2 | 1 | No |
| YP_009820823.1 | hypothetical protein HOV27_gp138 | 16 | 1 | Yes |
| YP_009820766.1 | hypothetical protein HOV27_gp081 | 2 | 1 | No |
| YP_009820875.1 | DNA Polymerase HOV27_gp244 | 2 | 1 | No |
| YP_009820762.1 | hypothetical protein HOV27_gp077 | 5 | 1 | No |

**Table S3.**
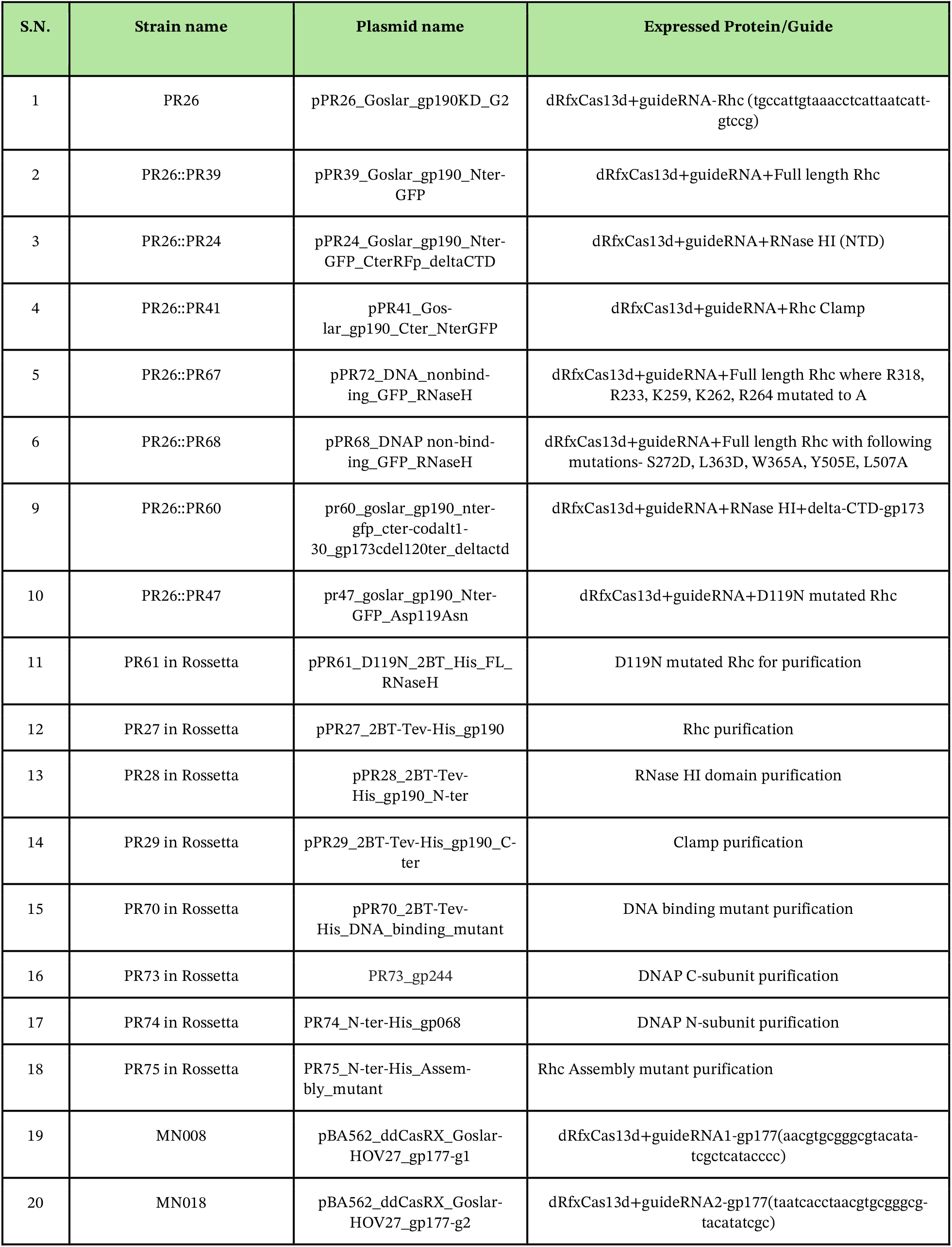
*E. coli MC1000 and Rossetta* strains were used in this study. GFP is “sfGFP” which is Addgene Plasmid #48885 pTD103luxI_GFP with mutations S2A, T65G, D190H and D197H. Plasmids with p15A-CmR backbone: PR26_Goslar_gp190KD_G2. Plasmids with pDSW206 backbone: PR39, PR24, PR41, PR60, PR47, PR67, PR68, and Plasmids with 2BT backbone are: PR27, PR28, PR29, PR61, PR70, PR73, PR74, PR75, P76.

**Table S4.**
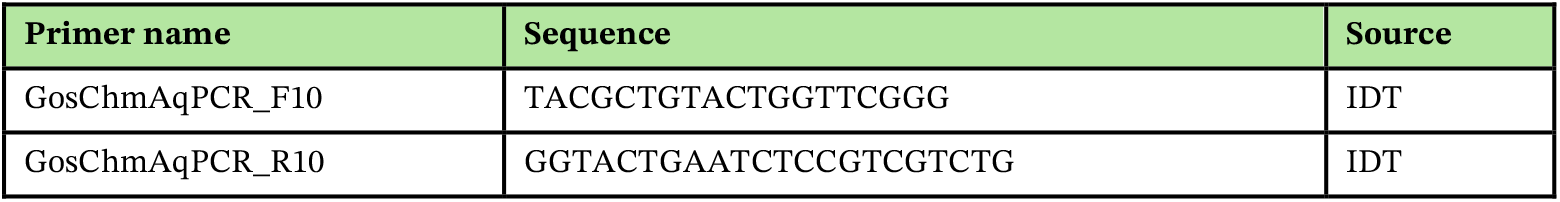
qPCR primers used in this study.

**Table S5. DESeq2 analysis of knockdown conditions compared with the control.**

**Table S6. RNA-seq quality control metrics and sequencing coverage across all samples.**

**Table S7. Comparative differential expression analysis of Rhc knockdown and ChmA knockdown relative to control.**

**Table S8.** Subset of RNA-seq differential expression results for Rhc KD versus NT control, filtered from Table S5.

| Gene ID | Base mean | log2 fold change | Adjusted P value | Status |
| --- | --- | --- | --- | --- |
| HOV27_gp246 | 58833.7 | 2.95 | 1.98E-106 | Upregulated |
| HOV27_gp214 | 13458.5 | 2.63 | 4.55E-81 | Upregulated |
| HOV27_gp176 | 7210.9 | 2.10 | 9.11E-46 | Upregulated |
| HOV27_gp144 | 4451.2 | 2.04 | 3.92E-52 | Upregulated |
| HOV27_gp022 | 435.3 | -7.43 | 7.32E-106 | Downregulated |
| HOV27_gp023 | 661.5 | -6.79 | 2.79E-171 | Downregulated |
| HOV27_gp217 | 1548.2 | -4.15 | 4.25E-56 | Downregulated |
| HOV27_gp093 | 289.0 | -4.01 | 2.73E-55 | Downregulated |
| HOV27_gp091 | 197.2 | -4.00 | 1.78E-24 | Downregulated |
| HOV27_gp056 | 599.3 | -3.72 | 4.01E-65 | Downregulated |
| HOV27_gp057 | 662.4 | -3.62 | 3.41E-36 | Downregulated |
| HOV27_gp203 | 841.8 | -3.62 | 1.45E-42 | Downregulated |
| HOV27_gp013 | 306.7 | -3.61 | 6.86E-30 | Downregulated |
| HOV27_gp024 | 450.5 | -3.57 | 8.24E-60 | Downregulated |
| HOV27_gp202 | 424.2 | -3.54 | 2.64E-45 | Downregulated |
| HOV27_gp092 | 1751.6 | -3.50 | 2.42E-51 | Downregulated |
| HOV27_gp076 | 276.2 | -3.43 | 2.11E-36 | Downregulated |
| HOV27_gp055 | 631.4 | -3.40 | 6.98E-56 | Downregulated |
| HOV27_gp079 | 388.5 | -3.28 | 5.77E-34 | Downregulated |
| HOV27_gp218 | 5635.3 | -3.28 | 2.51E-34 | Downregulated |
| HOV27_gp028 | 516.2 | -3.24 | 2.06E-51 | Downregulated |
| HOV27_gp030 | 696.6 | -3.23 | 1.42E-37 | Downregulated |
| HOV27_gp025 | 1827.2 | -3.23 | 1.6E-36 | Downregulated |
| HOV27_gp192 | 316.1 | -3.22 | 2.11E-36 | Downregulated |
| HOV27_gp204 | 3739.9 | -3.21 | 1.24E-36 | Downregulated |
| HOV27_gp026 | 2049.3 | -3.14 | 1.94E-37 | Downregulated |
| HOV27_gp027 | 623.0 | -3.11 | 7.18E-40 | Downregulated |
| HOV27_gp029 | 2272.4 | -3.10 | 4.88E-46 | Downregulated |
| HOV27_gp036 | 1517.1 | -3.03 | 8.42E-44 | Downregulated |
| HOV27_gp063 | 1710.3 | -3.03 | 7.48E-29 | Downregulated |
| HOV27_gp224 | 473.0 | -3.01 | 6.72E-41 | Downregulated |
| HOV27_gp062 | 603.2 | -2.95 | 4.9E-28 | Downregulated |
| HOV27_gp052 | 131.4 | -2.93 | 4.22E-17 | Downregulated |
| HOV27_gp090 | 1828.3 | -2.93 | 6.45E-31 | Downregulated |
| HOV27_gp189 | 527.0 | -2.92 | 1.16E-41 | Downregulated |
| HOV27_gp035 | 1350.0 | -2.90 | 2.36E-44 | Downregulated |
| HOV27_gp161 | 1926.1 | -2.74 | 1.76E-58 | Downregulated |
| HOV27_gp084 | 263.7 | -2.73 | 3.95E-21 | Downregulated |
| HOV27_gp033 | 891.5 | -2.71 | 1.93E-27 | Downregulated |
| HOV27_gp160 | 929.2 | -2.70 | 7.74E-32 | Downregulated |
| HOV27_gp219 | 658.3 | -2.68 | 1.09E-36 | Downregulated |
| HOV27_gp188 | 1018.6 | -2.67 | 8.1E-34 | Downregulated |
| HOV27_gp031 | 986.2 | -2.63 | 3.43E-32 | Downregulated |
| HOV27_gp086 | 1376.8 | -2.62 | 8.01E-30 | Downregulated |
| HOV27_gp205 | 756.0 | -2.61 | 8.14E-30 | Downregulated |
| HOV27_gp048 | 753.8 | -2.61 | 2.15E-39 | Downregulated |
| HOV27_gp061 | 1305.2 | -2.56 | 1.06E-30 | Downregulated |
| HOV27_gp032 | 977.9 | -2.55 | 4.26E-31 | Downregulated |
| HOV27_gp085 | 409.4 | -2.54 | 1.84E-26 | Downregulated |
| HOV27_gp087 | 366.0 | -2.54 | 1.66E-22 | Downregulated |
| HOV27_gp034 | 1208.2 | -2.53 | 4.05E-33 | Downregulated |
| HOV27_gp049 | 468.5 | -2.50 | 1.01E-29 | Downregulated |
| HOV27_gp187 | 461.0 | -2.43 | 6.42E-19 | Downregulated |
| HOV27_gp041 | 11605.9 | -2.28 | 4.56E-23 | Downregulated |
| HOV27_gp054 | 727.7 | -2.25 | 3.78E-28 | Downregulated |
| HOV27_gp088 | 1032.8 | -2.24 | 1.73E-14 | Downregulated |
| HOV27_gp051 | 873.4 | -2.13 | 6.2E-20 | Downregulated |
| HOV27_gp220 | 1725.4 | -2.12 | 1.01E-38 | Downregulated |
| HOV27_gp089 | 499.8 | -2.07 | 6.26E-20 | Downregulated |
| HOV27_gp060 | 1719.5 | -2.04 | 2.57E-17 | Downregulated |
| HOV27_gp058 | 6611.3 | -2.00 | 2.51E-21 | Downregulated |

**Table S9. Rhc_dependent_transcriptional_blocks.** Rhc_B1-Rhc_B16 denote 16 putative discrete genomic blocks of coordinately downregulated, adjacent genes following Rhc depletion. Block size indicates the number of genes within each block (2-5 genes), and position in block indicates gene order in the 5′→3′ transcriptional direction. Block start/end and length define the genomic coordinates and span of each affected region. baseMean represents mean normalized expression, log2FoldChange the expression change upon Rhc depletion, and pvalue/padj the statistical significance before and after multiple-testing correction, respectively.

**Table S10.** RNA-seq Rhc KD::WT* vs NT control.

| Gene ID | Base mean | log2 fold change | Adjusted P value | Status |
| --- | --- | --- | --- | --- |
| HOV27_gp059 | 1813.0 | 2.85 | 5.24E-32 | Upregulated |
| HOV27_gp042 | 1676.9 | 2.81 | 4.53E-28 | Upregulated |
| HOV27_gp058 | 6611.3 | 2.52 | 1.25E-32 | Upregulated |
| HOV27_gp041 | 11605.9 | 2.38 | 7.38E-25 | Upregulated |
| HOV27_gp190 | 17902.7 | 2.26 | 6.48E-08 | Upregulated |
| HOV27_gp218 | 5635.3 | 2.15 | 1.81E-15 | Upregulated |
| HOV27_gp157 | 3445.7 | 2.01 | 1.28E-73 | Upregulated |
| HOV27_gp182 | 1902.5 | -2.26 | 3.56E-69 | Downregulated |
| HOV27_gp156 | 675.2 | -2.25 | 1.18E-35 | Downregulated |
| HOV27_gp183 | 2223.3 | -2.21 | 1.24E-71 | Downregulated |
| HOV27_gp071 | 1775.2 | -2.12 | 9.36E-55 | Downregulated |
| HOV27_gp227 | 696.3 | -2.10 | 1.2E-34 | Downregulated |
| HOV27_gp075 | 1231.1 | -2.10 | 2.15E-41 | Downregulated |
| HOV27_gp070 | 1955.0 | -2.06 | 1.99E-66 | Downregulated |
| HOV27_gp225 | 1781.1 | -2.03 | 2.32E-41 | Downregulated |

